# SARS-CoV-2 Restructures the Host Chromatin Architecture

**DOI:** 10.1101/2021.07.20.453146

**Authors:** Ruoyu Wang, Joo-Hyung Lee, Feng Xiong, Jieun Kim, Lana Al Hasani, Xiaoyi Yuan, Pooja Shivshankar, Joanna Krakowiak, Chuangye Qi, Yanyu Wang, Holger K. Eltzschig, Wenbo Li

**Author notes:** Correspondence: Wenbo Li, Ph.D. MSB 6.161, Fannin Street, Houston, Texas, 77030, USA. These authors contributed equally (Ruoyu Wang, Joo-Hyung Lee).

## Abstract

SARS-CoV-2 has made >190-million infections worldwide, thus it is pivotal to understand the viral impacts on host cells. Many viruses can significantly alter host chromatin^1^, but such roles of SARS-CoV-2 are largely unknown. Here, we characterized the three-dimensional (3D) genome architecture and epigenome landscapes in human cells after SARS-CoV-2 infection, revealing remarkable restructuring of host chromatin architecture. High-resolution Hi-C 3.0 uncovered widespread A compartmental weakening and A-B mixing, together with a global reduction of intra-TAD chromatin contacts. The cohesin complex, a central organizer of the 3D genome, was significantly depleted from intra-TAD regions, supporting that SARS-CoV-2 disrupts cohesin loop extrusion. Calibrated ChIP-Seq verified chromatin restructuring by SARS-CoV-2 that is particularly manifested by a pervasive reduction of euchromatin modifications. Built on the rewired 3D genome/epigenome maps, a modified activity-by-contact model^2^ highlights the transcriptional weakening of antiviral interferon response genes or virus sensors (e.g., *DDX58*) incurred by SARS-CoV-2. In contrast, pro-inflammatory genes (e.g. *IL-6*) high in severe infections were uniquely regulated by augmented H3K4me3 at their promoters. These findings illustrate how SARS-CoV-2 rewires host chromatin architecture to confer immunological gene deregulation, laying a foundation to characterize the long-term epigenomic impacts of this virus.

## MAIN

The COVID-19 pandemic poses an unprecedented challenge to humankind. Mechanistic understanding of how SARS-CoV-2 impacts cellular functions is critical to solving this challenge. Control of host chromatin has been used by other viruses to antagonize host defense or to exert long-term influences ^1,3^, which, however, is largely unknown for SARS-CoV-2.

The folding of mammalian chromatin in three-dimension (3D) is an outstanding but incompletely understood process in biology^4^. Several layers of architectures are now observed that together influence critical nuclear processes such as gene transcription, replication, recombination, or DNA damage repair^4–6^. These architectures include A/B compartments, Topological Associating Domains (TADs), and chromatin loops^5, 7^ (**Extended Data Fig. 1a,b**). A/B compartments are suggested to be formed, at least in part, via homotypic attractions between chromatin regions of similar epigenetic features^7, 8^. For TADs and loops, CCCTC-binding factor (CTCF) and the cohesin complex are two main regulators^6^. Growing evidence supports that cohesin acts via a “loop extrusion” process by which it actively translocates on chromatin to create DNA contacts inside TADs, until it encounters specific CTCF sites and was blocked from further extrusion^6^ (**Extended Data Fig. 1a,b**). These two mechanisms can crosstalk – the formation of TADs by loop extrusion appears to antagonize compartmentalization^6,7,9,10^. How 3D chromatin architectures are rewired in pathological conditions remains poorly understood.

### Widespread restructuring of the host 3D genome

To explore how SARS-CoV-2 may impact the host 3D genome, we conducted viral infection in A549 cells (of human alveolar epithelial cell origin) expressing Angiotensin-converting enzyme 2 (ACE2) receptor (A549-ACE2) (**Fig. 1a**, **Extended Data Fig. 1c**), and first generated ribo-depleted total RNA-Seq at two time-points post infection (see data summary in **Extended Data Table 1**). A 24-hour infection (multiplicity of infection or MOI: 0.1) resulted in ∼90% of RNA-Seq reads aligned to the viral genome, indicating a high level of infection (**Extended Data Fig. 1d**), in agreement with previous observations^11^. Shorter term infection for 6-hr resulted in ∼20% RNA-Seq reads attributed to the virus genome, suggesting insufficient viral infection/replication (**Extended Data Fig. 1d**)^11^. We therefore focused on cells 24hr-post-infection (24hpi) to characterize the viral impacts on the host chromatin. Consistent with RNA-Seq, immunofluorescence of the spike protein verified a high ratio of viral infection at 24hpi (**Extended Data Fig. 1e**).

**Fig. 1.**
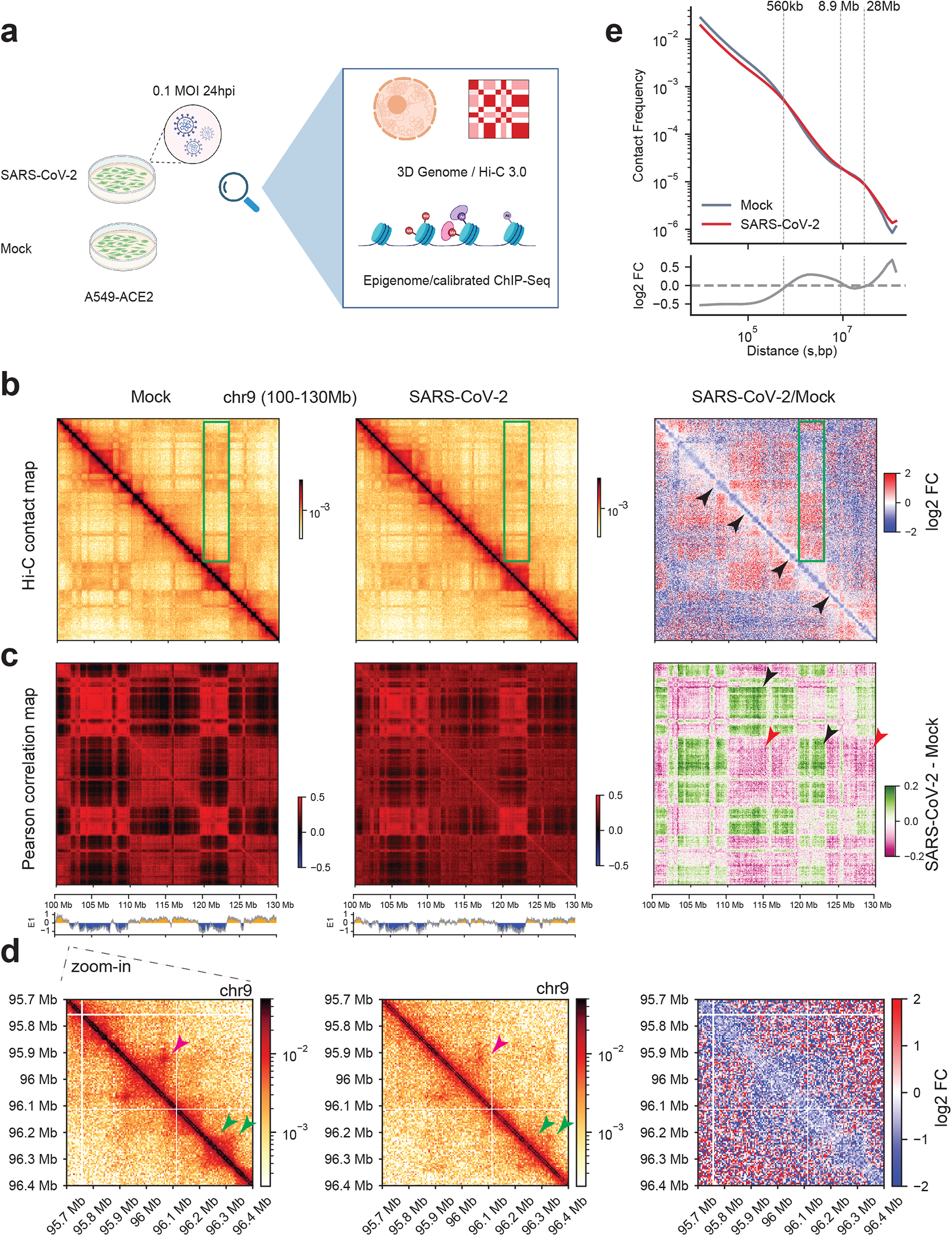
SARS-CoV-2 significantly restructured the host 3D genome. **a.** A diagram illustrating the experimental design of this work. **b.** Snapshots of normalized contact matrices of Hi-C 3.0 at an example region (chr9:100-130Mb, hg19) in Mock or SARS-CoV-2 infected conditions. Log2 fold change of Hi-C contact is shown to the right. Black arrowheads point to reduced short-distance interactions along the diagonal. Green boxes show altered compartmentalization. **c.** Snapshots of Pearson correlation matrices of Hi-C 3.0 in the same region as panel b. Changes of the correlation matrices after infection are shown to the right. Arrowheads point to regions with altered Pearson correlation matrices and are changed A-B (black) or A-A (red) compartmental interactions. **d.** Zoom-in Hi-C snapshots of a 700kb region in panel b,c (chr9:95.7-96.4Mb, hg19). Pink and green arrowheads show changed or unchanged domains and dot-shaped chromatin loops. **e.** (Top) P(s) curve showing the relationship between contact frequency (P) and distance (s) of both mock (grey) and SARS-CoV-2 (red) Hi-C datasets. (Bottom) Log2 fold changes of contact frequency ranked by distances, with dotted lines marking the crossing points of the two curves.

We employed Hi-C 3.0^12^, a recent version of *in situ* Hi-C that revised cross-linking procedure and included two restriction enzymes, which can detect high-resolution 3D chromatin architecture at both long and short distances. We generated biological replicates in A549-ACE2 cells at 24hpi or in mock-infected cells (Mock), and deeply sequenced the Hi-C 3.0 libraries to ∼2.6 billion read pairs for mock and infected conditions (**Extended Data Table 1**), generating ∼630-770 millions unique contact pairs for each condition. There is high concordance between Hi-C 3.0 replicates as demonstrated by the stratum-adjusted correlation coefficients (SCC)^13^ (**Extended Data Fig. 2a,b**), and an example region is shown in **Extended Data Fig. 2c**. We thus combined the replicates for the analysis below, and still refer to these data as Hi-C.

Hi-C revealed a strikingly widespread alteration of the 3D genome in SARS-CoV-2 infected cells. Evidently, the near-diagonal value of the contact matrix that denotes short range chromatin interactions was weakened globally, as exemplified by a 30Mb region on chromosome 9 (**Fig. 1b**, black arrowheads). Whereas the contacts far away from the diagonal, which are long-distance chromatin interactions, were often deregulated (increased or decreased for different regions, green box) (**Fig. 1b**). Pearson correlation map of Hi-C interactions consistently revealed these changes (**Fig. 1c**), which also suggests alteration of chromatin compartmentalization (see below). Zoom-in view of an example region of ∼0.7Mb showed that chromatin domains (“rectangles” with high-frequency cis interactions, **Extended Data Fig. 1a,b**) are often weakened (**Fig. 1d**), whereas chromatin loops (“dot” structures off the diagonal, **Extended Data Fig. 1a,b**) are deregulated (**Fig. 1d**). A P(s) curve shows the frequency (P) of intra-chromosomal interaction ranked by genomic distance (s) (**Fig. 1e**), demonstrating that SARS-CoV-2 elicited a global reduction of short distance chromatin contact (< 560kb), a moderate increase of mid-to-long distance interactions (∼560k-8.9Mb), and enhanced interactions for far-separated regions (>28Mb). Intriguingly, at the inter-chromosomal levels, trans-chromosomal contacts were generally increased by viral infection, as shown by the fold changes of pairwise interactions between any two chromosomes, or by calculating the ratios of trans-versus-cis chromosomal contacts in Hi-C data (**Extended Data Fig. 2d,e**). The enhancement of both inter-chromosomal interactions and the extremely long-distance (>28Mb) intra-chromosomal interactions (**Fig. 1e**) suggest changes in chromatin compartmentalization (see below). At such large scales, inter- and intra-chromosomal interactions can display some shared properties^9, 14^.

### Defective compartmentalization, A weakening, and A-B mixing

Principal component analysis (PCA) of Hi-C maps can divide 3D genomes into A/B compartments, which largely overlap the euchromatin and heterochromatin, respectively^7, 15^. Regions in each compartment preferentially interact within the same compartment^7, 15^ (**Extended Data Fig. 1a,b**). Analyzing a 100kb-binned Hi-C contact matrix, we found significant defects of chromatin compartmentalization in SARS-CoV-2 infected cells (**Fig. 2a**). Overall, compartmental scores (E1) exhibited a general reduction in virus infected cells (i.e., moving below the diagonal, **Fig. 2a**), suggesting a pervasive weakening of A compartment and/or A-to-B switching. By using a E1 score change of 0.2 as cutoff (see **Methods**^16^), we found that ∼30% of genomic regions exhibited compartmental weakening or switching after viral infection (**Extended Data Fig. 3a**). Consistent with the general trend of E1 reduction, compartmental changes commonly display features of weakened A (e.g. A to weaker A, or A to B) or strengthening of B compartment (B to stronger B). Among these, A-to-weaker-A change is the most common (∼18% of all genomic bins, **Fig. 2a** and **Extended Data Fig. 3a**). Example regions are shown in **Fig. 2b**.

**Fig. 2.**
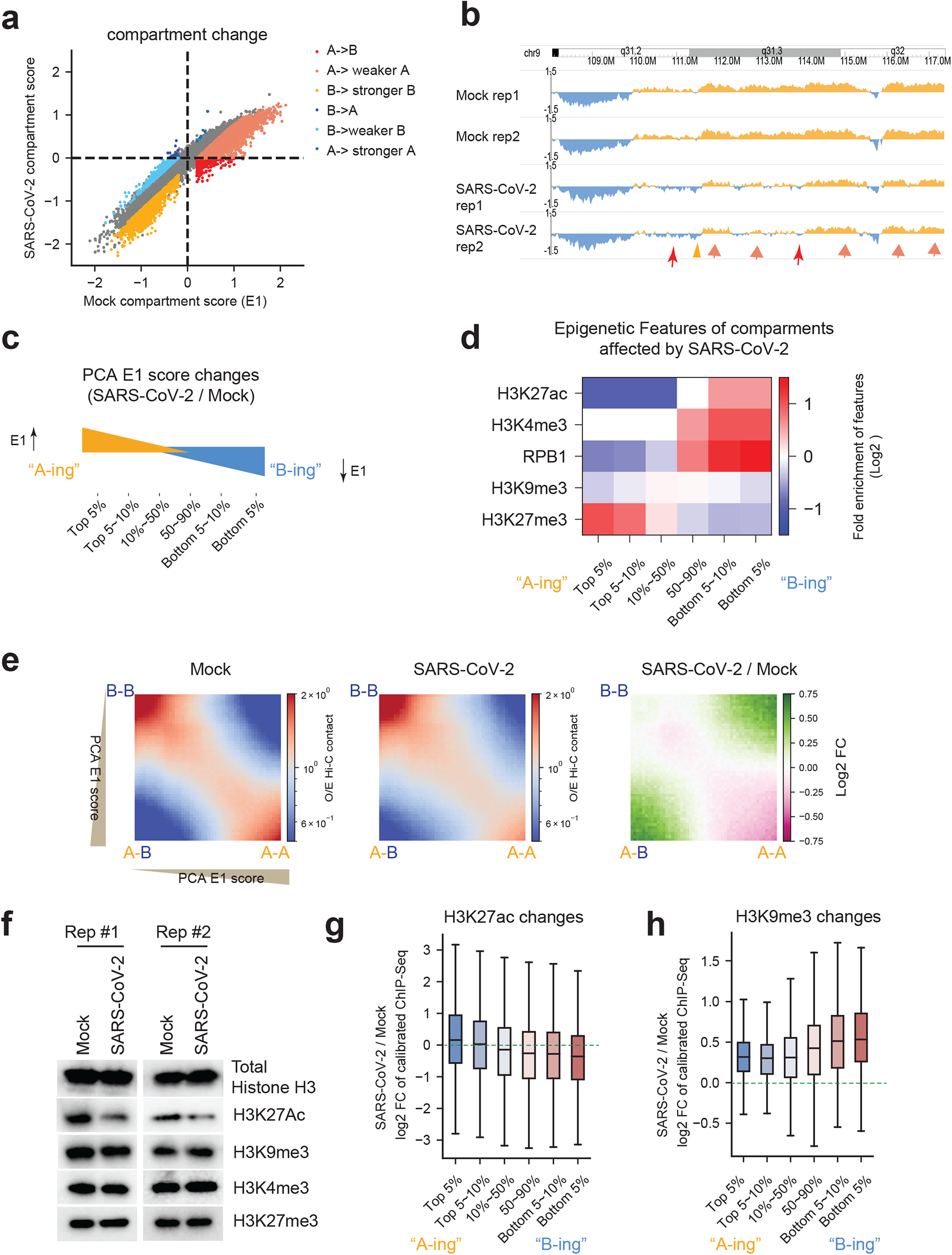
Compartmental A weakening, A-B mixing and epigenome reprogramming by SARS-CoV-2. **a.** A scatter plot showing the compartmental scores (E1 scores) of genome-wide 100kb genomic bins in mock or SARS-CoV-2 infected cells. Six categories of changes based on E1 changes were colored-coded. **b.** A snapshot of compartmental score (E1) tracks at an example region (chr9: 108Mb-118Mb). Rep1/Rep2 are replicates. Orange or blue color indicates A compartment (E1 score >0) or B compartment (E1 score < 0), respectively. Arrows and arrowheads exemplify compartmental changes in panel a (with matched colors). **c.** A diagram shows the categories of genomic bins with changes of A/B compartmental scores, and were referred to as “A-ing” (E1 scores increase) or “B-ing” (E1 scores reduce) after virus infection. **d.** Heatmaps showing the enrichment of active or repressive histone marks or RNA Pol2 (RPB1) on the six categories of genomic bins defined in panel c. Scale indicates log2-transformed enrichment of these features (see **Methods**). **e.** Saddle plots showing the compartmentalization between genomic regions ranked by their PCA E1 scores (all genomic regions were divided into 50 bins in total, see **Methods**). Therefore, for example, A-A homotypic interactions are in the right lower part of the plot; A-B interactions are in the right upper and left lower parts. Differential compartmental interactions are shown on the right as log2 fold changes (SARS-CoV-2/Mock). **f.** Western blots showing the abundances of total histone H3 or several modifications in Mock and SARS-CoV-2 infected (24hpi) cells. **g.** Boxplots showing the log2 fold changes of calibrated ChIP-Seq signals of H3K27ac and H3K9me3 for the six categories of genomic bins with varying compartmental changes (as in panel c). The boxplot centre lines represent medians; box limits indicate the 25th and 75th percentiles; and whiskers extend 1.5 times the interquartile range (IQR) from the 25th and 75th percentiles.

We evaluated the epigenetic features of the regions prone to compartmental changes incurred by the virus. By ranking each genomic bin based on its E1 score change, we sorted them into six categories (**Fig. 2c**). For those showing E1 increase, we dubbed them “A-ing” bins and those showing decrease “B-ing”. We then examined the enrichment of both active and repressive histone marks on these six categories of bins. Genomic regions originally harbored higher active chromatin marks known to feature A compartment (such as H3K27ac) were found to become more B-like (“B-ing” bins) after viral infection; whereas the genomic regions that were originally higher in repressive histone marks known to feature B compartment (particularly H3K27me3) become more A-like (“A-ing” bins) by SARS-CoV-2 (**Fig. 2d**). RNA polymerase II (Pol2) exhibits similar enrichment as that of H3K27ac mark (**Fig. 2d**). These results suggest that both the originally A or B compartments were losing their identity, indicating defective chromatin compartmentalization.

Indeed, defects of chromatin compartmentalization can be clearly detected in the example region shown in **Fig. 1c**, in which the originally strong separation between A and B compartmental regions become “fuzzier”. Inter-compartmental interactions formed between regions of A and B increased, whereas those formed homotypically within A or B reduced (**Fig. 1c**). A saddle plot demonstrated that such changes are global after virus infection (**Fig. 2e**, see **Methods**^8, 10^). There is a strong reduction of A-A homotypic interactions accompanied by gain of A-B mixing, while B-B interactions were not obviously changed (**Fig. 2e**). Similarly, by calculating another compartmentalization metric, the sliding correlation scores (SC score^9^, **Extended Data Fig. 3b**, see **Methods**), we found that SC scores were not affected at boundaries where A/B compartments transition to the other, but were globally decreased for regions inside the same compartments. This result reinforces the conclusion that SARS-CoV-2 weakened host chromatin compartmentalization. Increased A-B mixing (and hence weakened compartmentalization) can also be seen at very large scales in between two chromosomes, e.g. chromosomes 17 and 18, where originally well-separated A-A/B-B homotypic interactions were significantly compromised but A-B interactions globally enhanced (**Extended Data Fig. 3c**).

### Viral reprogramming of host epigenome and H3K27ac reduction

A-compartment in general enriches active histone modifications, while B-compartment repressive and heterochromatin marks^15^. To mechanistically understand compartmentalization changes, we systematically profiled epigenomic landscapes in Mock or SARS-CoV-2 infected cells by ChIP-Seq of active and repressive histone modifications (H3K4me3, H3K27ac, H3K9me3, and H3K27me3) (**Fig. 1a**). Western blots showed that most of these modifications remain unaltered, but surprisingly, active histone mark H3K27ac displayed a consistent and significant reduction after SARS-CoV-2 infection (**Fig. 2f**). We therefore conducted spike-in calibrated ChIP-Seq to precisely quantify epigenomic changes (see **Methods**^17^, **Extended Data Fig. 4a**). By calculating the ratio of ChIP-Seq reads aligned to the human versus to the mouse genome (i.e., spike-in), we consistently observed a reduction of H3K27ac on the host chromatin by ∼40-45% (**Extended Data Fig. 4b,d**), which agrees with the changes in western blots (**Fig. 2f**). There is no overall change of another active mark H3K4me3 (**Extended Data Fig. 4c,e**), but there are moderate gains of repressive histone marks, particularly the H3K9me3, after infection (**Extended Data Fig. 4c,f,g**).

The epigenome reprogramming after SARS-CoV-2 infection resonates with the pervasive decrease of A-A compartmental interactions and weakening of A compartment in Hi-C data (**Fig. 2a,e**). We examined histone mark changes on the six categories of genomic regions that bear differential compartmental changes (**Fig. 2c**), finding that the most weakened A compartment (“B-ing”) regions are associated with stronger reduction of H3K27ac (**Fig. 2g**), whereas heterochromatin mark H3K9me3 displays the opposite trend (**Fig. 2h**). An example is shown in **Extended Data Fig. 4h**, where H3K27ac reduced and H3K9me3 increased, correlating with compartmental changes: weakened A-A contacts and increased A-B inter-compartmental mixing. Because attractions between homotypic chromatin regions were suggested to be an important basis for compartmentalization^7,9,10^, our results support the notion that SARS-CoV-2 disrupted host chromatin compartmentalization, at least in part, via reprogramming chromatin modifications.

### Pervasive weakening of intra-TAD interactions

We then examined chromatin architectures at finer scales (10kb∼1Mb), namely TADs and chromatin loops^5–7^. A very pronounced phenomenon as shown by example regions in **Fig. 1d** and **Fig. 3a** is that cis-interactions within TADs were significantly reduced by virus infection, whereas the contacts beyond TADs (out of the rectangle) are unchanged or increased. Examining this phenomenon genome-widely, we calculated the insulation scores (IS)^18^ on our Hi-C maps and identified 4,094 TADs (see **Methods**). Aggregation Domain Analyses (ADA) of these TADs verified the significant weakening of intra-TAD cis-interactions by SARS-CoV-2, which accompanies unchanged or increased cis-interactions outside of TADs (**Fig. 3b**). Quantification of all intra-TAD contacts showed dramatic reduction (**Fig. 3c**). Interestingly, weakened intra-TAD contacts were not accompanied by severe loss of TAD identities, i.e. the boundaries of TADs were largely unchanged (**Fig. 3a,b** middle panel). Indeed, insulation scores on TAD boundaries were only mildly affected by SARS-CoV-2 (**Fig. 3d**).

**Fig. 3.**
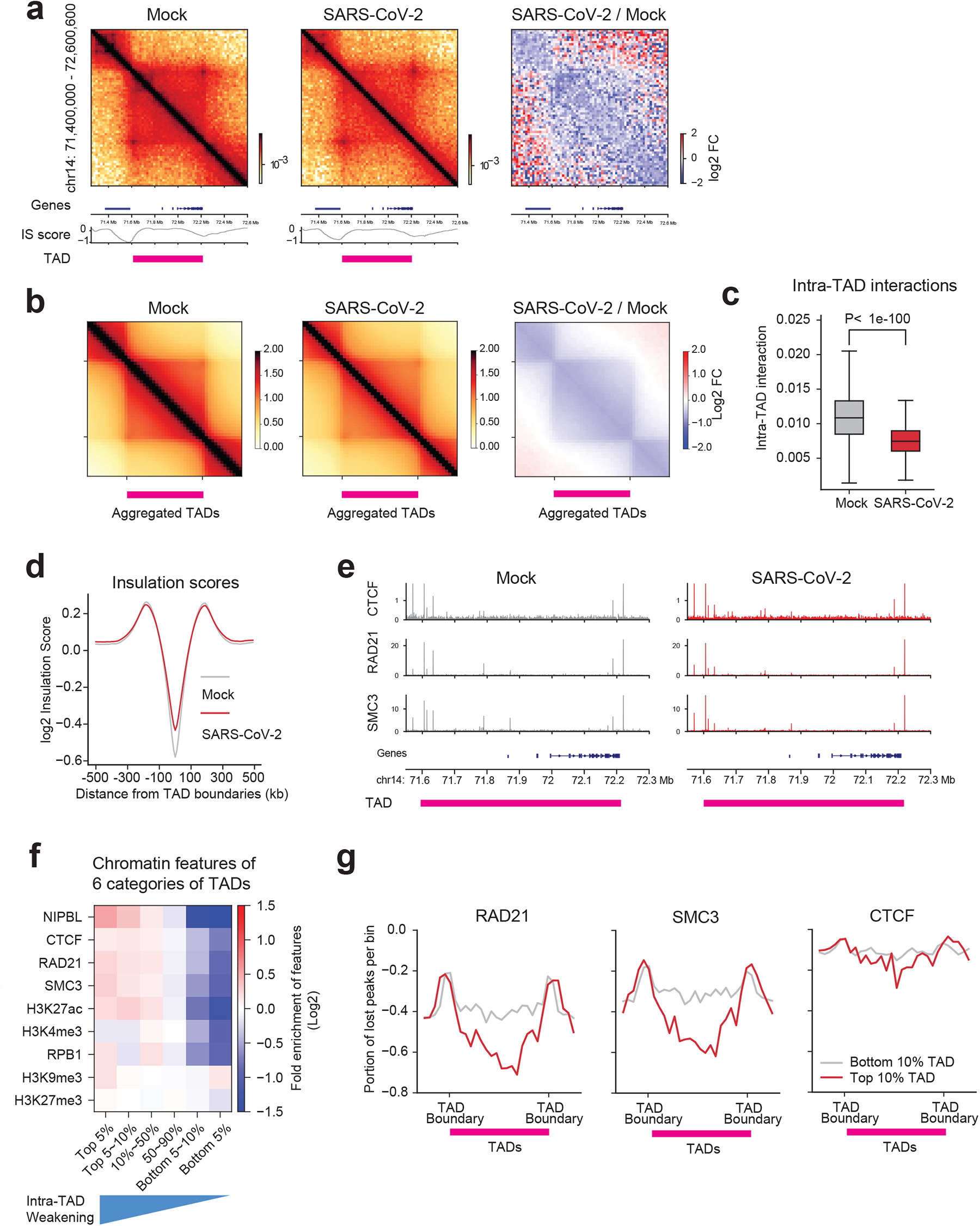
SARS-CoV-2 pervasively weakens intra-TAD contacts and cohesin loading. **a.** Snapshots of Hi-C contact matrices at the indicated region (hg19) in Mock and SARS-CoV-2 conditions. Also shown are the log2 fold changes of the Hi-C interactions (right panel), RefSeq gene tracks (bottom), the Insulation score (IS score, bottom), and TAD location (pink bars). Genomic coordinate is indicated on the left. **b.** Aggregated domain analyses (ADA) plots showing genome-wide reduction of intra-TAD interactions after SARS-CoV-2 infection. Right: log2 fold changes of signals. TAD location is denoted by a pink bar. The plots include additional 0.5xTAD on each side for quantification. **c.** A boxplot showing intra-TAD interactions in Mock and SARS-CoV-2 infected conditions. The boxplot centre lines represent medians of intra-TAD interaction from 4,094 TADs; box limits indicate the 25th and 75th percentiles; and whiskers extend 1.5 times the interquartile range. P-value: Mann-whitney U test. **d.** A profile plot of insulation scores calculated based on Mock (grey) and SARS-CoV-2 (red) infected Hi-C at all TAD boundaries. **e.** Snapshots of CTCF, RAD21, SMC3 ChIP-Seq tracks of both mock (left) and SARS-CoV-2 (right) conditions at an example region also shown in the panel a (chr14:71.5-72.3Mb, hg19). **f.** A heatmap showing the enrichment of cohesin/CTCF or epigenetic marks in six categories of TADs ranked by their quantitative reduction of intra-TAD interactions (also see **Extended Data Fig. 6a** and **Methods**). **g.** A meta-profile showing the portions of ChIP-Seq peaks reduced by SARS-CoV-2 for RAD21, SMC3 and CTCF, and the relative positioning of these changes to TADs (see **Methods**). Line plots show the distribution of lost peaks over a region centered on the midpoint of each TAD (0.25xTAD on each side out of TAD). Red lines indicate lost peaks belong to the top 10% weakened TADs, grey lines indicate the 10% least weakened TADs. The y-axis indicates the portion of lost peaks versus total peaks (e.g. -0.6 indicates 60% peaks in that bin were lost).

### Cohesin depletion from intra-TAD regions

To understand the drastic diminishing of intra-TAD interactions, we examined the chromatin binding of CTCF and cohesin, the main organizers of TADs^5–7^ by calibrated ChIP-Seq. These factors were not affected by 24hpi infection at protein levels (**Extended Data Fig. 5a**). In accord, spike-in calibration of ChIP-seq showed no significant global change of their chromatin binding (**Extended Data Figs. 4a**, **5b**). Peak calling for each factor results in ∼40,000∼60,000 peaks (**Extended Data Fig. 5c,e,g**). Importantly, for two cohesin subunits, 40.4% (27,152/67,140) of RAD21 sites and 31.8% (20,837/65,379) of SMC3 sites were significantly reduced after SARS-CoV-2 infection (**Extended Data Fig. 5c,d,e,f** see **Methods**), indicating a dramatic depletion of cohesin from chromatin. In contrast, only a small percentage of their binding sizes were gained, i.e., 2.2% (1,510/67,140) for RAD21 sites and 3.1% (2,034/65,379) for SMC3 (**Extended Data Fig. 5c,e**). Moderate changes of CTCF binding were observed, with 10.6% (4,853/45,530) sites lost and 10.0% (4,555/45,530) sites gained due to SARS-CoV-2 (**Extended Data Fig. 5g,h**). These changes can be seen in an example TAD region (**Fig. 3e**) whose Hi-C map was shown in **Fig. 3a**. We divided all TADs into six categories based on their sensitivity to SARS-CoV-2 infection (i.e., quantitative reduction of intra-TAD contacts, **Extended Data Fig. 6a**), and examined the chromatin features of these categories. Notably, the more virus-sensitive the TADs are (i.e., more weakening of intra-TAD contacts), the higher enrichment of cohesin and NIPBL they bear (**Fig. 3f**), suggesting that intra-TAD weakening is due to cohesin depletion in these TADs. Indeed, as compared to the least weakened TADs, the top 10% most virus-weakened TADs are associated with a more dramatic loss of cohesin from the intra-TADs regions (**Fig. 3g**). In addition, cohesin loss was more dramatic at the intra-TAD regions, although was detectable at the TAD boundaries (**Fig. 3e,g**). As comparisons, CTCF was minimally impacted either at TAD boundaries or intra-TADs (**Fig. 3e,g**), indicating that SARS-CoV-2 preferentially disrupts cohesin action inside TADs, but largely leaves intact the TAD structures.

Besides cohesin, virus-sensitive TADs were originally more enriched in active chromatin mark H3K27ac and RNA Pol2 (RPB1), but possess comparable heterochromatin marks as compared to other TADs (**Fig. 3f**). We thus examined the epigenetic changes of these TADs after infection, finding that for H3K27ac, whereas its level was globally reduced (**Fig. 2f**), the reduction was consistent among all categories of TADs (**Extended Data Fig. 6b**). In contrast, H3K9me3, but not another heterochromatin mark H3K27me3, was gained significantly more in the most weakened TADs (**Extended Data Fig. 6c,d**). This unique correlation suggests that H3K9me3 increase may play a role in cohesin depletion and intra-TAD weakening caused by SARS-CoV-2.

### SARS-CoV-2 mildly impacted dot-shaped chromatin loops

Dot-shaped chromatin loop is a prominent feature of Hi-C data that often forms between convergent CTCF sites^7, 15^. The definition and functions of chromatin loops can be debatable^7^, for example, enhancer-promoter contacts may be defined as loops by other work or methods but often do not appear as dots in Hi-C^5, 7^. Some work defined dot-shaped loops in Hi-C largely as structural loops that may not be regulatory for gene transcription^19^. In this study, we refer to dot-shaped structures as chromatin loops, and define enhancer-promoter contacts using Hi-C interaction strength (see **Methods**). Deep Hi-C 3.0 data permitted us to detect 11,926 loops in Mock or SARS-CoV-2 infected cells at 5∼10 kb resolution by HICCUPS, which outnumbers loops often detectable by conventional *in situ* Hi-C^9, 12^. Some example regions are shown below in **Fig. 4b** and **Extended Data Fig. 8d,e**. At a global scale, aggregation peak analyses (APA) showed that chromatin loops were not overtly affected by viral infection (**Extended Data Fig. 7a**). By quantitative changes (Hi-C contacts change FDR<0.1, see **Methods**), we found that 2.96% (353/11,926) loops were weakened by 24hpi, whereas 4.70% (560/11,926) loops gained strength (**Extended Data Fig. 7b,c**). The weakened loops are mostly short-range loops (median size 150kb), while interestingly the gained ones are much longer (median size 417.5kb, (**Extended Data Fig. 7d**). This is consistent with the P(s) curve (**Fig. 1e**) that long distance chromatin interactions were enhanced by virus infection. As compared to virus-strengthened loops, the anchors of weakened short loops showed more dramatic cohesin depletion (**Extended Data Fig. 7e**), and are more preferentially located intra-TADs than on boundaries (**Extended Data Fig. 7f,g**). These results are consistent with the pervasive reduction of intra-TAD interactions (**Fig. 3b,c**), suggesting that the virus-weakened short-range loops are a consequence of defective cohesin loop extrusion inside TADs. That the majority of loops are unaffected is consistent with that TAD boundaries were largely spared (**Fig. 3b,d**), where cohesin binding was mildly affected (**Fig. 3e,g**).

**Fig. 4.**
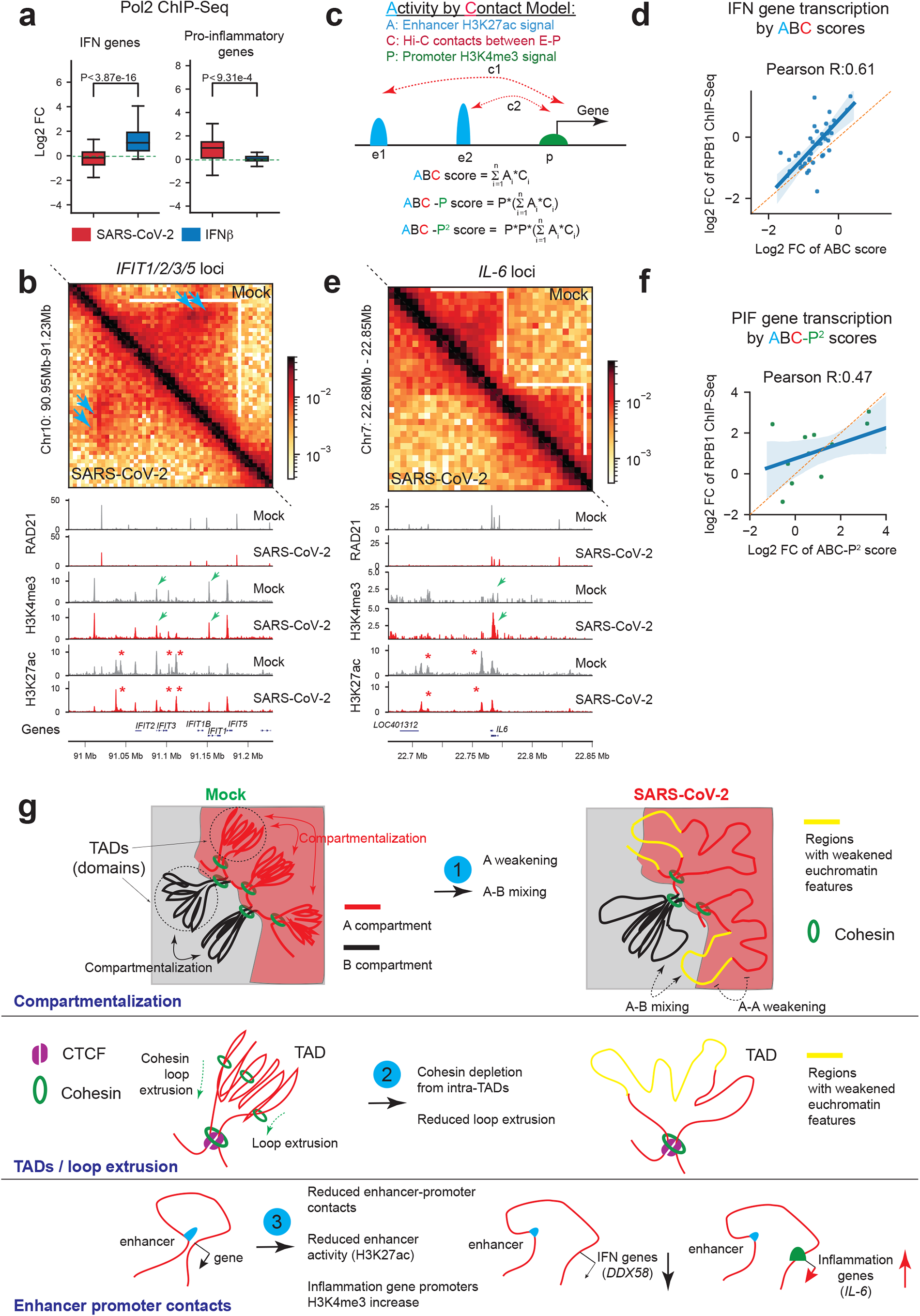
Chromatin restructuring underlies dichotomic transcriptional deregulation of interferon response genes and inflammation genes. **a.** Boxplots showing transcriptional deregulation of key interferon response gene (IFN) and pro-inflammatory genes after SARS-CoV-2 infection or IFN-beta treatment (1000u, 6-hr), as shown by fold changes of RNA Pol2 (RPB1) ChIP-Seq. The boxplot centre lines represent medians; box limits indicate the 25th and 75th percentiles; and whiskers extend 1.5 times the interquartile range (IQR) from the 25th and 75th percentiles. P-value: Mann-whitney U test. **b.** Snapshots of Hi-C contact matrices and calibrated ChIP-Seq tracks for indicated factors at the *IFIT* gene cluster. Top right: Mock; lower left: SARS-CoV-2. Blue arrows point to reduced dot-shaped loops. White lines mark the TAD, with intra-TAD interactions weakened throughout. Red asterisks show virus-reduced H3K27ac peaks. Green arrows show unchanged H3K4me3 peaks on *IFIT1/3* promoters. **c.** A diagram shows the method of Activity-by-contact (ABC) to model the transcriptional outputs of a gene based on considering its enhancer activity and enhancer-promoter contacts; ABC-P or ABC-P^2^ are revised ABC algorithms that also consider promoter strength (see **Methods**). **d.** A scatter plot showing the correlation between the ABC scores (x-axis) and true transcriptional changes of IFN genes caused by SARS-CoV-2 (y-axis, RPB1 ChIP-Seq). A high Pearson’s correlation coefficient is achieved and labeled. A liner regression fitted line and its 95% confidence interval (shaded area) are also shown. **e.** Similar to panel b, the snapshots of Hi-C matrices and ChIP-Seq tracks at the *IL-6* locus. Top right: Mock; lower left: SARS-CoV-2. White lines mark the TADs, with intra-TAD interactions weakened throughout. Red asterisks show reduced H3K27ac peaks. Green arrows show enhanced H3K4me3 on *IL-6* promoter. **f.** A scatter plot showing the correlation between the ABC-P^2^ scores (x-axis) and true transcriptional changes of pro-inflammatory genes by SARS-CoV-2 (y-axis, RPB1 ChIP-Seq). Pearson’s correlation coefficient is shown. A liner regression fitted line and its 95% confidence interval are shown. **g.** A proposed model that summarizes our findings of the chromatin restructuring by SARS-CoV-2 at scales of both 3D genome and 1D epigenome. They are categorized in compartments, TADs/loop extrusion and enhancer-promoter contacts (see **Extended Data Fig. 1a,b** for additional information). These changes can explain transcriptional deregulation of interferon response and inflammation genes that confer immunological phenotypes in COVID-19 patients.

### Restructured chromatin architecture explains immuno-pathological changes

COVID-19 patients with severe symptoms often show two key immuno-pathological features: a delayed/weakened innate immune antiviral response (i.e., type-I interferon response) together with an exacerbated production of pro-inflammatory cytokines (e.g., IL-6)^20^. Our maps of 3D genome and epigenome provided an opportunity to mechanistically understand the deregulation of these genes incurred by the virus. Our RNA Pol2 (RPB1) ChIP-Seq and RNA-Seq in 24hpi SARS-CoV-2 infected cells recapitulated these two immuno-pathological features, indicating that immunological gene alteration occurs transcriptionally (Pol2 in **Fig. 4a**, RNA-Seq in **Extended Data Fig. 8a**). As compared to interferon-beta (IFN-b) stimulus, SARS-CoV-2 infection elicited very limited activation of interfero<u>n</u> response genes (hereafter IFN genes) but strong increases of <u>p</u>roinflammatory genes (hereafter PIF genes)(**Fig. 4a**). Key examples include *IFIT1/2/3/5*, *DDX58* (a.k.a. *RIG-I*) or *IFIH1* (a.k.a. *MDA5*) for IFN genes, and *IL-6* or *CXCL8* for the PIF genes (**Extended Data Fig. 8b,c**). Inspection of IFN gene loci revealed remarkable changes of chromatin architecture by viral infection: 1), the dot-shaped loops were often diminished and the chromatin contacts (which include most enhancer-promoter contacts) throughout the hosting TAD were reduced; 2), cohesin occupancy within the hosting TADs was decreased; 3), the active mark H3K27ac displayed significant reductions at many putative enhancers near IFN genes (**Fig. 4b**, **Extended Data Fig. 8d,e**). Both enhancer-promoter contacts and enhancer activities may contribute to gene transcriptional outputs^2^ (**Extended Data Fig. 1a,b**), we thus modified and applied the activity-by-contact (ABC)^2^ algorithm to model how chromatin architecture and epigenome changes underlie virus rewiring of host transcription (**Fig. 4c**, also see **Methods**). Combining alteration of enhancer activity and enhancer-promoter contact strength correctly modeled the weakened transcriptional outputs of IFN genes (Pearson’s correlation coefficient R=0.61, **Fig. 4d**). In contrast, poor prediction was achieved if only one of the two features was considered (contact only: R=0.13 or enhancer activity only: 0.33; **Extended Data Fig. 8f,g**). We experimentally tested the functions of these enhancers by CRISPRi (dCas9-KRAB-MeCP2 and gRNAs) in two IFN gene loci where they were weakened by SARS-CoV-2, finding that IFN genes were indeed transcriptionally inhibited from responding to poly(I:C), a synthetic dsRNA simulates virus infection (**Extended Data Fig. 8h,i**). These results demonstrated that weakened 3D chromatin contacts and enhancer activity together shaped the transcriptional inhibition of IFN genes after SARS-CoV-2 infection.

However, for PIF genes, while ABC scores correlated with their changes, their true transcriptional levels after SARS-CoV-2 infection were often several-fold higher than ABC scores (**Extended Data Fig. 9a**, examples in **Extended Data Fig. 9b**). Indeed, as shown for *IL-6* and *CXCL8* (IL-8) loci, coding for key clinically detrimental cytokines, enhancer activities (H3K27ac) and intra-TAD contacts were reduced for these genes (**Fig. 4e**, **Extended Data Fig. 9c**), albeit they were strongly up-regulated by virus infection (**Extended Data Fig. 9b**). We therefore re-examined our epigenome datasets at the PIF gene loci, finding a unique and dramatic gain of H3K4me3 mark at their promoters (**Fig. 4e**, **Extended Data Fig. 9c**), which did not take place on IFN genes (**Fig. 4b**, **Extended Data Fig. 8d,e**). At a genome-wide scale, H3K4me3 exhibited relatively limited changes by SARS-CoV-2: 5.8% (1,843/31,761) sites showed increases and 3.5% (1,104/31,761) sites decreases (**Extended Data Fig. 9d**). Interestingly, the genes close to gained H3K4me3 sites are specifically enriched for TNF-alpha pathway, TGF-beta signaling, and inflammatory response (**Extended Data Fig. 9e**). Thus, we revised the ABC algorithm by including H3K4me3 changes at PIF gene promoters (**Fig. 4c**), and this revised ABC-P^2^ score better modeled their strong transcriptional induction (**Extended Data Fig. 9b**), **Fig. 4f**, see **Methods**). Intriguingly, virus-augmented promoters (with H3K4me3 increases) display motif enrichment of specific transcriptional factors, such as IRF1/2 or Jun/AP1 (**Extended Data Fig. 9f**). These results together suggest that inflammatory genes are induced by SARS-CoV-2 through a unique mechanism to augment their promoter strength (H3K4me3), even when their enhancer and enhancer-promoter contacts were weakened.

## DISCUSSION

Here, we mapped high-resolution landscapes of the 3D genome and epigenome in cells of human lung epithelial origin after acute SARS-CoV-2 infection, and our results revealed dramatic viral restructuring of the host chromatin. Hi-C 3.0 data uncovered significant defects of chromatin compartmentalization and TAD structures (**Fig. 4g**). These are manifested at the compartment levels as wide-spread A weakening and A-B mixing, and at the TAD levels as global reduction of intra-TAD chromatin contacts. These 3D genome changes, to our knowledge, represent a unique and previously unappreciated rewiring of the mammalian 3D genome in pathological settings.

Notably, the cohesin complex was depleted, in a pervasive but also selective manner, from intra-TAD regions by SARS-CoV-2. Such changes of cohesin not only provide a molecular explanation to the weakening of intra-TAD chromatin contacts, but also support a notion that cohesin extrusion defects inside TADs released these chromatin regions to engage in long-distance chromatin associations (**Fig. 4g**). Indeed, virus infected chromatin displayed a higher frequency of extremely long distance intra-chromosomal as well as inter-chromosomal interactions (**Fig. 1e** and **Extended Data Fig. 2d**). Gains of long distance chromatin interactions have been observed after genetic depletion of cohesin subunits from human or mouse cells, supporting that active loop extrusion prevents chromatin mixing due to homotypic attraction^6,9,10^. Two critical features highlight that SARS-CoV-2 infection disrupts both the epigenome and cohesin loop extrusion, resulting in 3D genome changes in a manner more complex than genetic depletion of cohesin: 1), the augmented compartmental interactions after SARS-CoV-2 infection are mostly formed inter-compartmentally (i.e., between A-B), whereas those after cohesin depletion were mostly formed intra-compartmentally (e.g., between A-A); 2), SARS-CoV-2 infection significantly ablates cohesin from intra-TAD regions, but affects them mildly at TAD boundaries. The first feature can be explained by a global weakening of euchromatin modification after virus infection. Indeed, virus-elicited decreases of active mark H3K27ac and increases of heterochromatin mark H3K9me3 correlate with the weakening of A compartment. In contrast, histone modifications were largely unchanged by genetic depletion of cohesin^9^. We propose that weakened euchromatin after SARS-CoV-2 infection on one hand reduced A compartment strength and A-A interactions, and on the other blurred the distinction between A and B compartments, contributing to wide-spread A-B mixing. This is consistent with the notion that homotypic chromatin attractions, once disrupted, can compromise compartmentalization^7, 8^. For the second feature, the mechanisms underlying preferential cohesin depletion from intra-TAD regions remain to be determined. We found that TADs most sensitive to SARS-CoV-2 are originally of higher enrichment of cohesin loader NIPBL, suggesting a plausible scenario that this virus utilizes specific mechanisms to perturb cohesin loading in these TADs. Alternatively, the cohesin release process may also be perturbed, which affects the loading-release equilibrium of cohesin to allow its preferential loss inside TADs^6^.

An important insight from our work is that architectural restructuring of host chromatin underlies the dichotomic transcriptional changes of immunological genes seen in COVID-19 pathology: weakened IFN responses accompanying increased pro-inflammatory gene expression^20^. Weakening of enhancer-promoter contacts together with reduced enhancer activity correlates well with inhibited IFN gene transcription, and was validated by CRISPRi experiments. These changes take place at critical disease loci, including genes coding for key RNA virus sensors, *DDX58* (coding for RIG-I), whose inhibition is required for successful infection^21^. Unexpectedly, we found that SARS-CoV-2 directly upregulates the promoter activities of pro-inflammatory genes, suggesting unappreciated regulators on these promoters to confer inflammatory phenotypes in COVID-19 patients^20^. Further studies are warranted to understand such promoter-centered PIF transcriptional activation, which may offer therapeutic targets to relieve lethal inflammation in patients.

Epigenetic alteration is known to exert long term effects in affecting gene expression or cellular phenotypes^22, 23^. Understanding the viral alteration of host chromatin not only provides new knowledge and strategies to fight SARS-CoV-2, but also paves way for further studies of the epigenetic disturbance in patients suffering long COVID^24^.

## Contribution

W.L. conceived the project. Wet lab: J.-H.L., and R.W. did most of the experiments, with help from F.X., L.A.H., and J.K. J.Kim, P.S, X.Y., J.Y.W. and H.K.E. generated SARS-CoV-2 virus and conducted human cell infections. Dry lab: R.W. Manuscript: W.L., R.W. and J.-H.L. wrote the manuscript, and received inputs from all authors.

## Acknowledgements

W.L. is a Cancer Prevention and Research Institute of Texas (CPRIT) Scholar. This work is supported by funding from the University of Texas McGovern Medical School, NIH ‘‘4D Nucleome’’ program (U01HL156059), NCI (K22CA204468), NIGMS (R21GM132778, R01GM136922), CPRIT (RR160083, RP180734), Welch foundation (AU-2000-20190330) and the John S. Dunn foundation to W.L, as well as from NIH (R01HL154720, R01DK122796, R01DK109574, R01HL133900) and Department of Defense (W81XWH2110032) to H.K.E.. J.-H.L. is a recipient of a UTHealth Innovation for Cancer Prevention Research Training Program Post-doctoral Fellowship (RP160015). R.W. is a recipient of a John and Rebekah Harper Fellowship in Biomedical Sciences (to MD Anderson and UTHealth Graduate School). Most of our next generation sequencing work was conducted with UTHealth Cancer Genomics Core, which received funding from CPRIT (RP180734).

## Contact for Reagent and Resource Sharing

All requests for information, reagents and resources should be directed to the Lead Contact: Wenbo Li, Ph.D..

## Data availability statement

Data generated in this study have been deposited to NCBI GEO (GSE179184, https://www.ncbi.nlm.nih.gov/geo/query/acc.cgi?acc=GSE179184).

## Code availability statement

Analyses of Hi-C, ChIP-Seq and RNA-Seq in this study were performed by using published softwares or codes as described in the Methods part.

## Materials and Methods

### Cell culture

Human lung adenocarcinoma cells A549 expressing human ACE2 (A549-ACE2, #NR53821) was acquired from BEI Resources (Manassas, VA). They were maintained in DMEM/F-12 (1:1, Corning) medium supplemented with 10% FBS (GeneDepot) and blasticidin (100uM). Normal A549 cells were purchased from ATCC (CCL-185) and were cultured in DMEM/F-12 (1:1, Corning) supplemented with 10% FBS. 293T cells were from ATCC and were cultured in DMEM with 10% FBS. Vero-E6 cells were acquired from ATCC (CRL-1586). Mouse ESCs (F121-9) are a gift from David Gilbert lab (Florida State University) and were cultured following standard procedure of the 4D nucleome consortium (https://data.4dnucleome.org/biosources/4DNSRMG5APUM/). All these cells were cultured at 37°C with 5% CO2. Transfection of plasmids or siRNAs was performed using Lipofectamine 3000 or RNAiMAX (Life technologies) following manufacturer’s instructions. For CRISPRi experiments, in order to examine enhancer functions during cell responses to RNA virus, we introduced poly(I:C) (333ng/ml, Sigma, P9582) into A549 cells using lipofectamine 2000 and harvested the total cellular RNAs for gene expression experiments after 4 hours.

### SARS-CoV-2 infections in A549-ACE2 cells

SARS-CoV-2 isolate USA-WA1/2020 (NR-52281; BEI Resources, Manassas, VA) was employed to infect human A549-ACE2 cells (NR53821; BEI Resources). For viral infections, serum/antibiotics-free Eagle’s MEM medium supplemented with 1mM HEPES was used. Briefly, cells grown in 10-cm culture dishes at about 70-80% confluency were washed with the serum free medium, viral inoculum was added at 0.1 MOI for 1 hr. After that, non-adsorbed viral particles were gently aspirated out and the monolayers were replenished with 10% FBS containing MEM supplemented with 1mM HEPES. Infected cells were incubated at 37°C with 5% CO2 for 6hr or 24hr post-infection for experiments.

### Preparation of SARS-CoV-2 stock

The stock SARS-CoV-2 was propagated in Vero-E6 cells. Briefly, Vero-E6 cells were grown to 80% confluence in 10% FBS containing MEM medium supplemented with 1mM HEPES and 1X antibiotics and antimycotics. Prior to infection, Vero-E6 cells were washed once with PBS and the viral inoculum was added to the flask in the presence of 3 ml of serum-free and antibiotics-free MEM medium supplemented with 1mM HEPES, and incubated for 1 hour at 37°C with 5% CO2. At the end of incubation, non-adsorbed virus was aspirated out and cells were replenished with 25ml of MEM supplemented with 10% FBS and 1mM HEPES. Infected cells were incubated for 48 hours at 37°C with 5% CO2. At 80% of cell lysis, SARS-CoV-2 was harvested by detaching all the cells with a cell scraper and centrifuging at 300g for 3 minutes. Viral aliquots were stored in screw-cap vials at -80°C.

### Determination of plaque forming units (PFU/ ml stock)

For the determination of infectious viral titers, plaque assays were performed using Vero-E6 cells. Briefly, Vero-E6 grown in 6-well plates were infected with 12 serial dilutions (1:10) of the SARS-CoV-2 stock in serum/BSA/antibiotic-free MEM medium with 1mM HEPES for 1 hour at 37°C with 5% CO2. At the end of incubation, non-adsorbed viral particles were aspirated and the infected cells were layered upon with MEM medium containing 0.5% agarose, 2% BSA and 1mM HEPES, and incubated for 48 hours at 37°C with 5% CO2. Fixation was carried out using 3.75% buffered formaldehyde (in PBS) for 10 minutes. After aspirating formaldehyde, the agarose layers were gently removed. Infected cells were stained with 0.3% crystal violet for 5 minutes, followed by washing once with PBS. Plates were air-dried and visible infectious plaques were counted in each dilution to determine the plaque forming units/ ml of the stock.

### Lenti-viral Transduction and CRISPRi

We in-house generated a lentiviral construct expressing dCas9-KRAB-MeCP2 by PCR amplification of the dCas9-KRAB-MeCP2 (contains a domain of MeCP2) from pB-CAGGS-dCas9-KRAB-MeCP2 (Addgene 110824), and then insert it to the pLenti-EF1a-dCas9-VP64-2A-Blast backbone (Addgene 61425) to replace th dCas9-VP64. The gRNAs used in CRISPRi were cloned into the Addgene 61427 backbone using BsmBI enzyme. To generate lentivirus, 293T cells were transfected with the lentiviral transfer vector DNA, psPAX2 packaging and pMD2.G envelope plasmid DNA at a ratio of 4:3:1, respectively, by lipofectamine 2000. After 16 h, the culturing media was changed to fresh one, and the supernatants were collected twice at 48h to 72h post-transfection. The harvested lentiviral supernatants were filtered using 0.45 μm syringe filter (Fisher) and used to infect target A549 cells (polybrene was added at a final concentration of 8 μg/ml, Sigma). To infect A549 cells for CRISPRi, cells were first infected by a lentivirus expressing dCas9-KRAB-MeCP2 for 24 hours and selected with appropriate antibiotics (10 μg/ml blasticidin) for 7 days to generate a stable cell line. The stable cell line was then subjected to viral infection by individual gRNAs targeting each enhancer, and they were further selected by 100 μg/ml Zeocin for 4-7 days. These stable cells were then used for experiments. The gRNA cloning oligos are shown in **Extended Data Table 2**.

### RNA Extraction and RT-qPCR

RNA extraction of SARS-CoV-2 infected A549-ACE2 cells was performed by TRIzol (Thermo Fisher Scientific, 15596-026) following manufacturer’s instructions. In some other cases, RNAs of cells expressing CRISPRi or other transfection were performed using Quick-RNA Miniprep Kit (Zymo Research, 11-328). Reverse transcription was conducted by using Superscript™ IV first strand synthesis kit (Thermo Fisher, 18091050), and often the random hexamer primer was used to test the expression levels of target genes. Real-time qPCR (RT-qPCR) was performed using SsoAdvanced™ Universal SYBR® Green Supermix (Bio-Rad, 172-5274). Primer sequences used in this study can be found in **Supplementary Table 2**. Relative gene expression was normalized to internal control (18S RNA).

### Western blots

Cells were lysed in RIPA (50 mM Tris, pH 7.4, 150 mM NaCl, 1 mM EDTA, 0.1% SDS, 1% NP-40, 0.5% sodium deoxycholate) with cOmplete™ Mini Protease Inhibitor Cocktail (Roche, 11836153001) on ice for 30min. Lysates were sonicated in Qsonica 800R (25% amplitude, 3 min, 10 sec on 20 sec off interval) and centrifuged at 14,000rpm. The supernatants were mixed with 2x Laemmli Sample Buffer (Bio-Rad) and boiled at 95℃ for 10min. The boiled proteins were separated on 4% to 15% SDS-PAGE gradient gels and transferred to LF PVDF membrane (Bio-Rad, Cat #1620260). The membranes were blocked in 5% skim milk in TBST (20 mM-Tris, 150 mM NaCl, and 0.2% Tween-20 (w/v)) for an hour and then briefly washed in TBST twice. Then, the membranes were incubated in TBST with primary antibodies (GAPDH (Proteintech, 60004-1), RAD21 (Abcam, Ab992, Lot: GR214359-10), CTCF (Millipore, 07-729), SMC3 (Abcam, Ab9263, Lot:GR466-7), Total Histone H3 (Abcam, Ab1791, Lot:GR206754-1), H3K4me3 (Abcam, Ab8580, Lot: GR3264490-1), H3K9me3 (Abcam, Ab8898, Lot: GR164977-4), H3K27ac (Abcam, Ab4729, Lot: GR3357415-1), and H3K27me3 (Cell Signaling Technology, #9733S, Lot19)) at 4°C overnight. After washing 3 times in TBST, the blots were incubated in TBST with secondary antibody (Horseradish peroxidase (HRP)-conjugated antibody) for an hour. After 6 times of washing in TBST, the blots were developed in a Bio-Rad ChemiDoc™ Gel Imaging System.

### Immunofluorescence microscopy

Expression of spike protein of SARS-CoV-2 was measured by immunofluorescence microscopy. A549-ACE2 cells seeded on glass slides were infected with SARS-CoV-2 at a MOI of 0.1. 24hr post infection (24hpi), cells were fixed by 4% paraformaldehyde in phosphate buffered saline (PBS) for 1hr at room temperature. The coverslips were washed with 0.1% BSA in 1xPBS (Wash buffer) and blocked with 1% BSA with 0.3% Triton-X100 in PBST (Blocking buffer) for 45min at room temperature. Cells were incubated with 1:500 anti-SARS-CoV-2 Spike glycoprotein antibody (Abcam, Catalog#ab272504) diluted in blocking buffer for overnight at 4 degree. Subsequently, after washes, cells were incubated with 1:500 Alexa Fluor 594-conjugated-anti-rabbit-IgG (Jackson Immunoresearch, Catalog#111-585-144) diluted in blocking buffer for 1hr at room temperature, followed by incubation with 4,6-diamidino-2-phenylindole (Invitrogen, Catalog#D1306) for 5 minutes at room temperature. Coverslips were mounted in antifade mounting medium (Thermo scientific, Catalog#TA-030-FM) and the fluorescence images were recorded using a Leica confocal microscope.

### Hi-C 3.0

Hi-C 3.0 was performed based on a recent protocol^12^, which is largely modified based on in situ Hi-C^15^. Briefly, ∼5 million SARS-CoV-2 infected A549-ACE2 cells were washed once with PBS to remove debris and dead cells, trypsinized off the culture plates, and were cross-linked using 1% formaldehyde for 10mins at room temperature, quenched with 0.75M Tris-HCl pH 7.5 for 5mins. These cells were further cross-linked with 3mM disuccinimidyl glutarate (DSG) for 50mins, and again quenched with 0.75M Tris-HCl pH 7.5 for 5mins. Cross-linked cell pellets were washed with cold PBS, and then resuspended in 0.5 ml ice-cold Hi-C lysis buffer (10 mM Tris-HCl, pH 8.0; 10mM NaCl, 0.2% NP-40 and protease inhibitor cocktail), and rotated at 4°C for 30 mins. Nuclei were washed once with 0.5 ml ice-cold Hi-C lysis buffer. After pelleting down the nuclei, 100 μl of 0.5% SDS was used to resuspend and permeabilize the nuclei at 62 °C for 10 mins. Then 260 μl H20 and 50 μl 10% Triton-X100 were added to quench the SDS at 37 °C for 15 mins. Subsequently, enzyme digestion of chromatin was performed at 37 °C overnight by adding an additional 50 μl of 10X NEB buffer 2, 100U MboI (NEB, R0147M) and 100U DdeI (NEB, R0175L). After overnight incubation, the restriction enzyme was inactivated at 62 °C for 20 mins. To fill-in the DNA overhangs and add biotin, 35U DNA polymerase I (Klenow, NEB, M0210) together with 10 μl of 1mM biotin-dATP (Jeana Bioscience), 1μl 10mM dCTP/dGTP/dTTP were added and incubated at 37°C for 1 hour with rotation. Blunt end Hi-C DNA ligation was performed using 5000 U NEB T4 DNA ligase with 10X NEB T4 Ligase buffer with 10mM ATP, 90 μl 10% Triton X-100, 2.2 μl 50mg/ml BSA at room temperature for four hours with rotation. After ligation, nuclei were pelleted down and resuspended with 440 μl Hi-C nuclear lysis buffer (50mM Tris-HCl pH7.5, 10mM EDTA, 1% SDS and protease inhibitor cocktail), and further sheared using the parameter of 10/20 secs ON/OFF cycle, 25% Amp, 4 mins by a QSonica 800R sonicator. Around 10% of the sonicated chromatin was subjected to overnight decrosslinking at 65°C, protein K treatment, and DNA extraction. After DNA extraction, biotin labelled Hi-C 3.0 DNAs were purified by 20 μl Dynabeads® MyOne™ Streptavidin C1 beads (Thermo Fisher 65002). The biotinylated-DNA on C1 beads was used to perform on-beads library making with NEBNext® Ultra™ II DNA Library Prep Kit for Illumina (NEB, E7645L) following manufacturer’s instructions. The sequencing was done on a NextSeq 550 platform with PE40 mode.

### Chromatin immunoprecipitation (ChIP-Seq) and spike-in calibrated ChIP-Seq

ChIP-Seq was performed as previously described with minor modifications26. For most ChIP-Seqs in A549-ACE2 cells with Mock treatment or 24-hr SARS-CoV-2 infection, ∼5 to 10% of mouse ESCs (F121-9, a gift from David Gilbert) were added as spike-in controls before sonication with equal proportions to the two human cell samples (see **Extended Data Fig. 4a**). For cell cross-linking for ChIP-Seq, briefly, the cells were trypsinized in Trypsin-EDTA (or Accutase for mESCs). After centrifugation, the cells were crosslinked by 1% formaldehyde (FA) in PBS for 10 min. The fixation steps were stopped in a quenching solution (0.75M Tris-HCl pH 7.5) for 10min. After centrifugation of the cells, we extracted the nuclei first by buffer LB1 [50 mM HEPES-KOH (pH 7.5), 140 mM NaCl, 1 mM EDTA (pH 8.0), 10% (v/v) glycerol, 0.5% NP-40, 0.25% Triton X-100 and 1×cocktail protease inhibitor], and then by LB2 [10 mM Tris-HCl (pH 8.0), 200 mM NaCl, 1 mM EDTA (pH 8.0), 0.5 mM EGTA (pH 8.0) and 1×cocktail protease inhibitor]. After centrifuge, the nuclei were suspended in buffer LB3 [10 mM Tris-HCl (pH 8.0), 100 mM NaCl, 1 mM EDTA (pH 8.0), 0.5 mM EGTA (pH 8.0), 0.1% Na-deoxycholate, 0.5% N-lauroyl sarcosine and 1×cocktail protease inhibitor], and the chromatin was fragmented by using Q800R3 sonicator (QSonica) using conditions of 10 seconds ON, 20 seconds OFF for 7-9 mins (at 20% amplitube). Sheared chromatins were collected by centrifugation, and were incubated with appropriate antibodies (often 2-3μg) at 4°C overnight. The next morning, the antibody-protein-chromatin complex was retrieved by adding 25μl pre-washed Protein G Dynabeads (Thermo Fisher Scientific, 10004D). Immunoprecipitated chromatin DNA was de-crosslinked by 65°C heating overnight using elution buffer (1% SDS, 0.1M NaHCO3), and then treated by RNase A and proteinase K, which was finally purified by phenol chloroform. The DNAs were subjected to sequencing library construction using NEBNext® Ultra™ II DNA Library Prep Kit for Illumina (NEB, E7645L), and were deep sequenced on a NextSeq 550 platform using 40nt/40nt pair-ended mode. The antibodies used for ChIP-Seq include RNA Polymerase II (RPB1 N terminus, Cell Signaling Technology, #14958S, Lot4), RAD21 (Abcam, Ab992, Lot: GR214359-10), SMC3 (Abcam, Ab9263, Lot:GR466-7), CTCF (Millipore, 07-729), H3K4me3 (Abcam, Ab8580, Lot: GR3264490-1), H3K9me3 (Abcam, Ab8898, Lot: GR164977-4), H3K27ac (Abcam, Ab4729, Lot: GR3357415-1), H3K27me3 (Cell Signaling Technology, #9733S, Lot19) and HA (Abcam, Ab9110, Lot:GR3231414-3).

### Ribo-depleted total RNA sequencing (RNA-Seq)

Total RNAs from mock or virus infected A549-ACE2 cells were extracted by TRIzol, and 100-200ng of total RNAs were used for making strand-specific ribosome-RNA-depleted sequencing library by the NEB Ultra II Directional RNA library kit (E7760L) following manufacturer’s instructions. Libraries were sequenced on a NextSeq 550 using 40nt/40nt pair-ended mode.

### Bioinformatic analyses

#### Calibrated ChIP-Seq analyses

Sequencing reads were aligned to a concatenated genome of hg19 human genome assembly and mm9 mouse genome assembly with STAR v 2.7.027. Duplicated reads were removed, and only unique aligned reads will be considered for later visualization and quantification. The scaling factor was calculated as the ratio of the number of reads uniquely aligned to human chromosomes versus the number of reads aligned to mouse chromosomes (**Extended Data Fig. 4a**). Uniquely aligned human reads were extracted with samtools28, and normalized by the corresponding scaling factor with deeptools29. For RBP1 ChIP-Seq gene transcription quantification, hg19 RefSeq gene annotation coordinates were used. The peak-calling of most ChIP-Seq were performed with the parameters of -f BAM -q 0.01 in MACS2 30. For each peak, we considered it with a log2 fold change of calibrated ChIP-Seq signal greater than 1 or lower than one as gained peak or reduced peak. ChIP-Seq reads are summarized in Extended Data Table 1. A public NIPBL ChIP-Seq dataset was obtained from SRR3102878.

#### RNA-Seq analysis

RNA-Seq reads were aligned to the hg19 reference human genome or SARS-CoV-2 viral genome (NC_045512.2) with STAR v 2.7.027. The percentage of reads uniquely aligned to SARS-CoV-2 genome versus total reads was calculated to verify a high viral infection rate. For human gene quantification, only uniquely aligned reads mapped to the hg19 genome were kept for further analysis. Differential gene expression analyses were performed with EdgeR, and genes with |FC|>2, FDR<0.05 were considered as significantly differential expressed genes.

#### Hi-C 3.0 data processing

Hi-C 3.0 raw data was primarily processed with HiC-Pro^31^. The pair of reads were mapped to the human reference genome assembly hg19, and multi-mapped pairs, duplicated pairs, and other unvalid 3C pairs were filtered out following the standard procedure of HiC-Pro. All valid Hi-C pairs were merged between replicates (unless specified noted), and were further converted to Juicebox format^32^ or cooler format^33^ for visualization and further analyses. Hi-C contact matrices were normalized with *cooler balance* function^33^. Reads numbers of Hi-C are listed in **Extended Data Table 1**. The SCC correlation coefficients between two replicates were calculated to assess the reproducibility of Hi-C experiments^13^. The P(s) curve was calculated as a function between contact frequency (P) and genomic distances(s) (**Fig. 1e**). Only intra-chromosomal pairs (cis) were used to calculate P(s) curve.

#### A/B Compartment analyses

A/B nuclear compartments were identified based on decomposed eigenvectors (E1) from 20kb or 100kb Hi-C contact matrices using cooltools. A/B compartmental scores (E1) were corrected by GC densities in each bin. Saddle plot analyses were performed to measure the compartmentalization strength in a genome-wide scale using cooltools compute-saddle (similar to8,10). Briefly, we first sorted the rows and the columns in the order of increasing compartmental scores within observed/expected (O/E) contact maps based on the data in Mock cells. Then we aggregated the rows and the columns of the resulting matrix into 50 equally sized aggregate bins, and plotted the aggregated observed/expected Hi-C matrices as the “saddle” plots (**Fig. 2e**). In **Fig. 1c** and other few places, Pearson correlation Hi-C matrices were used to emphasize the compartmental checkerboard pattern. We first calculated the observed/expected Hi-C maps as OE matrices (bin size = 80,000 bp). Each value (i,j) in Pearson matrices indicates the Pearson correlation coefficient between the i-th column of OE matrices and the j-th column of OE matrices (bin size = 80,000bp). The sliding correlation score (**Extended Data Fig. 3b**) was obtained based on Pearson correlation matrices, and we largely follow a previous work9. Briefly, for each genomic bin i (bin size = 80,000bp), we calculated the Pearson correlation coefficient between the i-th column and i+1-th column of Pearson correlation matrices, as the sliding correlation score. This score indicates correlation for each region as compared to the neighboring region. Valleys of SC score imply strong differences in long-range contact pattern observed at a locus as compared to its neighboring loci, indicating a change in compartment. Compartmental domains are genomic regions with continuous positive or negative compartmental scores (E1), identified by applying HOMER tool (findHiCCompartments.pl) on E1 scores. For changes of compartmental strength (**Fig. 2a** and **Extended Data Fig. 3a**), the changes for each genomic region between Mock and SARS-CoV-2 samples were identified based on 100kb-binned compartmental scores (E1) of two Hi-C 3.0 replicates, largely following a previous study16. For each 100kb, a student’s t-test was first performed on Mock and SARS-CoV-2 compartmental scores (E1). Only the 100kb bins that have |delta E1| > 0.2 and P-value < 0.05 were considered as bins with changed compartmental strength. Different categories of compartment changes (in **Fig. 2a**) were defined as below (similar to16): A to stronger A: (Mock-E1 – SARS-CoV-2-E1) < -0.2, Mock-E1>0.2; B to A: (Mock E1 – SARS-CoV-2 E1) <-0.2, Mock-E1 < -0.2, SARS-CoV-2-E1>0; B to weaker B: (Mock-E1 – SARS-CoV-2 E1) < -0.2, Mock-E1 <-0.2, SARS-CoV-2-E1<0; B to stronger B: (Mock-E1 – SARS-CoV-2-E1)>0.2, Mock-E1 < -0.2; A to B: (Mock-E1 – SARS-CoV-2-E1)<-0.2, Mock-E1 > 0.2, SARS-CoV-2-E1<0; A to weaker A: (Mock-E1 – SARS-CoV-2-E1)<-0.2, Mock-E1 > 0.2, SARS-CoV-2-E1>0.

#### TADs and insulation scores

Hi-C 3.0 data were used to identify topologically associating domains (TADs) in A549-ACE2 cells following standard 4D Nucleome consortium protocol (github.com/4dn-dcic/docker-4dn-insulation-scores-and-boundaries-caller). First, insulation scores ^18^ and boundary strengths of each 10kb bin with a 200kb window size were measured to quantify the TAD insulation using cooltools (https://github.com/open2c/cooltools/blob/master/cooltools/cli/diamond_insulation.py). Then, we identified TAD boundaries respectively in Mock and SARS-CoV-2 infected samples by using a boundary score cutoff of 0.5. We further merged TAD boundaries identified in these two conditions, and compared insulation scores at merged TAD boundaries (**Fig. 3d**). Merged TAD coordinates were used to perform downstream analyses. For each TAD, we quantified its mean Hi-C contacts throughout the domain (excluding very short distant interactions <15kb), which is considered intra-TAD interaction used in the paper. Based on the log2 fold changes of intra-TAD mean Hi-C contacts (SARS-CoV-2 / Mock), we ranked all TADs into six categories (Top 5%, top 5∼10%, 10∼50%, 50∼90%, bottom 5∼10%, bottom 5%), and calculated different epigenomic features of these six categories. For histone modifications or chromatin regulatory factors that have sharp peaks in ChIP-Seq (like H3K27ac, H3K4me3, CTCF or cohesin subunits), we quantified the numbers of peaks or the numbers of gained or lost peaks in different TADs. For modifications or factors that have broad ChIP-Seq patterns (like H3K9me3 and H3K27me3), we quantified the calibrated ChIP-Seq reads throughout the TADs. The enrichments of these ChIP-seq signals were calculated by dividing the median or mean quantification inside these six categories by the genome-wide median or mean quantification.

#### Chromatin loop calling and enhancer-promoter contacts

For loop calling, we largely followed a recent 4DN benchmarking paper^12^. In brief, we used a reimplement of HICCUPS loop-calling tool, call-dots function inside cooltools (https://github.com/open2c/cooltools/blob/master/cooltools/cli/call_dots.py) to identify structural chromatin loops in different samples. We first called loops at 5kb and 10 kb resolution separately, then used the following strategy to merge 5kb and 10kb loops. 5kb loops called at both 10kb and 5kb resolution were first kept, all unique 10kb resolution loops were kept, and only unique 5kb loops that are smaller than 100kb were kept. Differential loops were identified by first quantifying the Hi-C raw contacts at 40kb resolution of each called loop, and then by performing DESeq2 differential analyses on these raw counts. We considered loops with a DESeq2 FDR <0.1 and a log2FC >0 or <0 as virus-strengthened or weakened chromatin loops. The APA (Aggregation Peak Analysis) was performed by superimposing observed/expected Hi-C matrices on merged loops with the coolpuppy tool^34^.

#### Activity by Contact (ABC) score

ABC score calculation largely follows a previous study2 with modifications. For A score (enhancer activity) of a gene, we first identified all putative enhancers of this gene by selecting H3K27ac ChIP-Seq peaks located within 1Mb of the promoter. Then we quantified the calibrated H3K27ac ChIP-Seq signals on these putative enhancers (extended 150bp from MACS2 peaks) as A scores. The A-only quantification of enhancer activity for this gene will be the sum of the A scores for all putative enhancers. For C score (enhancer-promoter contact) between a gene and putative enhancers, we quantified the normalized Hi-C contacts formed in between the 5kb bins harboring the gene promoter and the putative enhancer. For the ABC score, we multiplied the A score of each enhancer by the C score, and generated the summation of these if multiple putative enhancers exist for a gene. P score of any gene was calculated as the calibrated H3K4me3 ChIP-Seq signal at its promoter region (+/-2.5kb from TSS) of a gene. For ABC-P or ABC-P2 scores, we multiplied the summed ABC score of a gene by its P score (promoter H3K4me3 signal) or by the square of its P score. The transcriptional changes of any gene were calculated based on the log2 fold change of RBP1 ChIP-Seq reads over the whole gene body (average of three ChIP-Seq replicates). Pearson correlation coefficient was used to measure the correlation between ABC score change and transcriptional change. The list of interferon response (IFN) genes was obtained from GSEA molecular signature databases (Interfero_Alpha_Response), and the list of pro-inflammatory (PIF) genes was manually curated based on recent literature11 studying immuno-pathology of SARS-CoV-2 infection (see Extended Data Table 3).

#### Statistics

qPCR data was analyzed by Prism and presented as mean±SD, which are indicated in figure legends. At least two biological replicates were conducted for RNA-Seq, ChIP-Seq or Hi-C sequencing. Student’s t-test (two-tailed) was commonly used to compare means between two qPCR groups; p < 0.05 was considered significant, and we labeled the p values with asterisks in each figure panel (*, p < 0.05; **, p < 0.01; ***: p < 0.001). Statistical analyses for sequencing data were performed with Python or R scripts. Key softwares or algorithms used in our analysis of sequencing data are listed in methods.

## Extended Data Figure legends

**Extended Data Figure 1.**
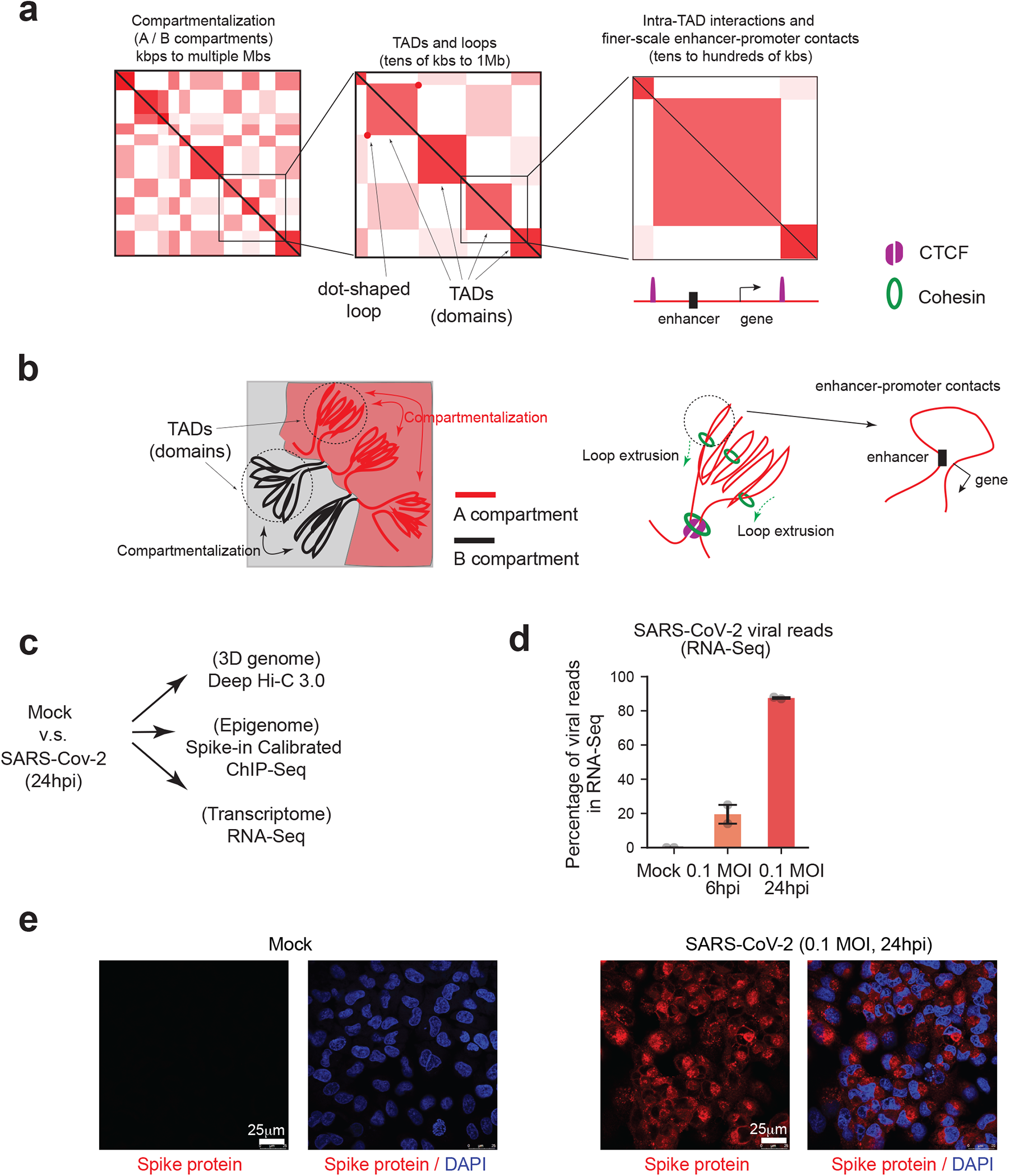
Overview of several layers of 3D chromatin architectures; overview and quality control of viral infection in our study. **a.** A diagram showing the typical contact map patterns in Hi-C (and Hi-C 3.0 or other modified Hi-C approaches) that define A/B compartments, Topologically Associating Domains (TADs), chromatin loops or intra-TAD interactions (which perhaps include most enhancer-promoter contacts). This is an overall summary of these structures, but the exact definition of some structures may be subjected to variable interpretation, and the terminology may not always be used consistently^5–7,15^. Often, A/B compartmentalization is illustrated by a checkerboard pattern of Hi-C contact matrices over large genomic sizes, indicating preferential interactions between genomic regions belonging to the same type of compartments (A: euchromatin and transcriptionally active; B: heterochromatin, transcriptionally inactive). TADs or chromatin domains are often characterized as a square or triangle-like structure on chromatin contact maps, reflecting a higher contact frequency between any regions inside the same TAD than with regions outside of the TAD. Intra-TAD enhancer-promoter contacts are considered to be facilitated by TADs, while TAD boundaries prevent aberrant interaction with regions outside of TADs. In Hi-C maps, the dot-shaped structures on the tip of domains suggests local enrichment of spatial interaction between a pair of two loci over nearby regions, and is regarded as a chromatin loop in this work. But loops may be subjected to other definitions in other studies. For example, enhancer-promoter contacts often do not appear as dot-shaped structures in Hi-C, but may be defined as loops by other work or other methods. Additional discussion, see^5, 7^. **b.** Cartoon diagrams describe A-A and B-B association preference within regions of similar epigenetic features, which compartmentalizes chromosomes into A and B (the left part of the diagram). The diagram in the middle depicts a current model of cohesin loop extrusion inside TADs that generated such structures. The right side shows a zoom-in view of a part of a TAD that harbors enhancer-promoter contact that may play roles in gene transcriptional regulation. **c.** A workflow showing the experimental design. **d.** A barplot showing the percentage of RNA-Seq reads mapped to SARS-CoV-2 genome in Mock, 6-hr post infection (6hpi, 0.1 MOI), and 24 hpi (0.1 MOI) conditions. Mean and standard deviation (error bar) were calculated based on two biological replicates of RNA-Seq. **e.** Confocal images showing immunofluorescence staining of DAPI (DNA, blue) and the Spike protein of SARS-CoV-2 (red) in Mock and 24hpi (0.1 MOI) infected A549-ACE2 cells. Scale bars are shown.

**Extended Data Figure 2.**
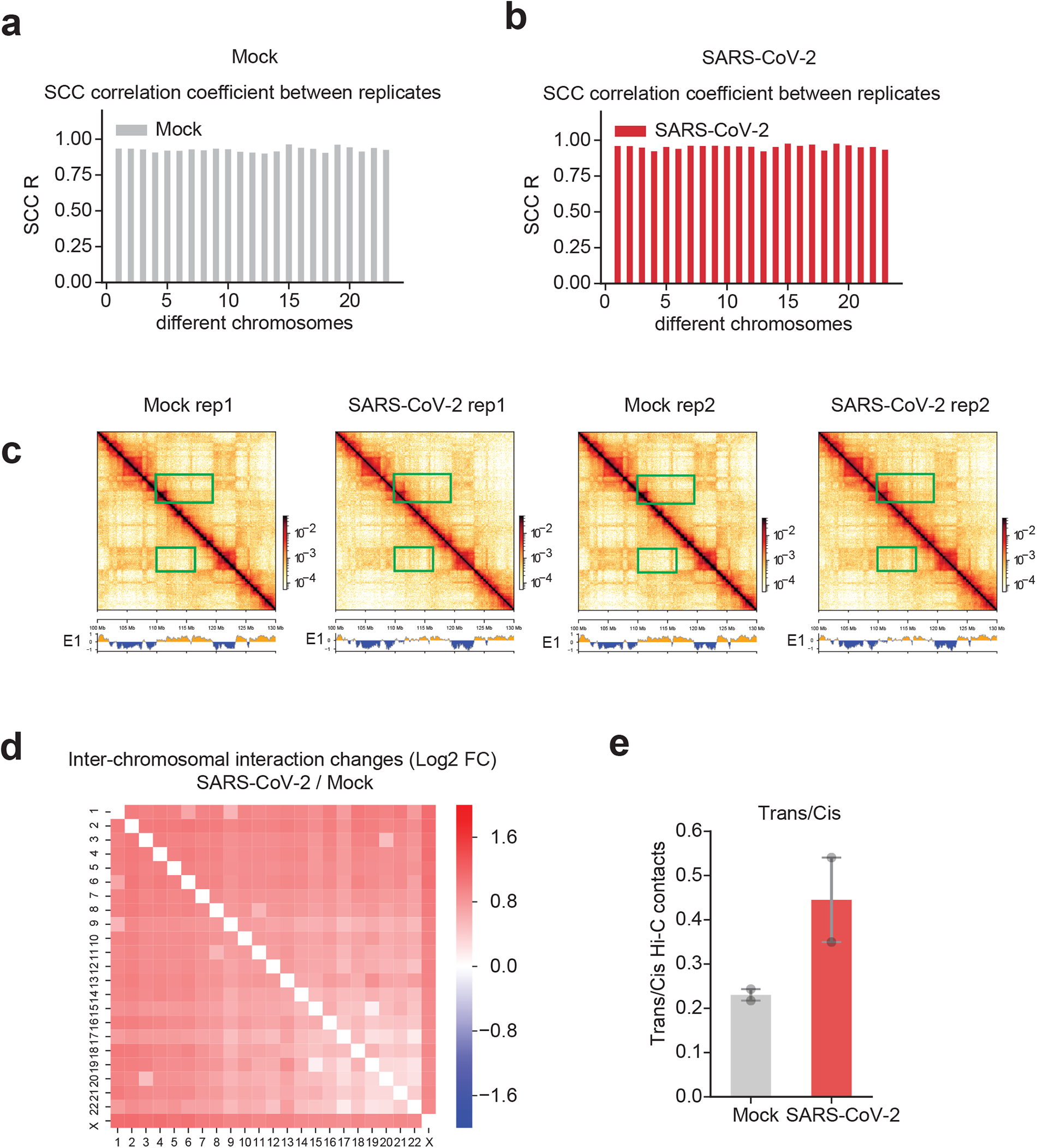
Replicates consistency of Hi-C 3.0, and the increase of trans-chromosomal interactions after SARS-CoV-2 infection. **a,b.** Barplots showing the SCC correlation coefficients^13^ between two Hi-C 3.0 replicates of Mock or SARS-CoV-2 conditions for different chromosomes. **c.** Snapshots showing two replicates of Hi-C contact matrices and compartmental score (E1) tracks in the same genomic region shown in **Fig. 1b**. The left two matrices show data of replicate 1 (rep1), and the right two matrices show replicate 2 (rep2). Green boxes show two regions with increased A-B compartmental mixing or weakened A compartment after virus infection. **d.** A heatmap shows the log2 fold change of inter-chromosomal interactions between two pairs of any two chromosomes (SARS-CoV-2/Mock). **e.** A barplot showing the trans/cis Hi-C contacts ratio in two replicates of Mock or SARS-CoV-2 infected cells. Trans contacts indicate chromatin interactions formed in between two different chromosomes. Mean and standard deviation (error bar) were calculated based on two biological replicates of Hi-C.

**Extended Data Figure 3.**
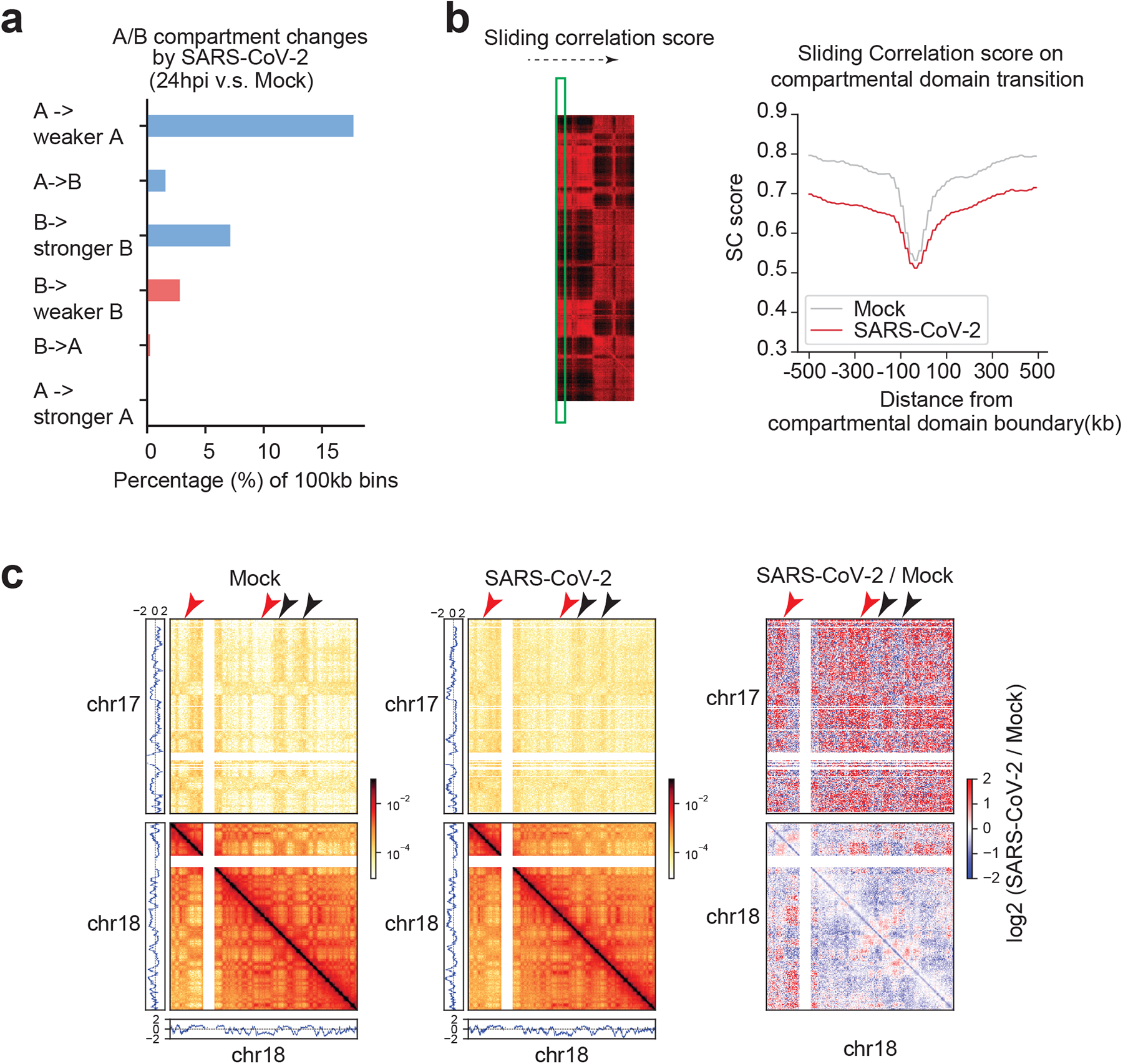
Additional data of A/B compartmental changes. **a.** A barplot showing the percentage of genomic bins (100kb bin size) that can be categorized into six groups based on their compartmental score changes (E1 value). These six categories are: A to weaker A, A to B, B to stronger B, B to weaker B, B to A, and A to stronger A. **b.** A diagram showing the basis of sliding correlation score to examine changes of compartmental interactions based on Pearson’s correlation matrices (see^9^, see **Methods**). On the right, a meta-profile plot of SC (sliding correlation) scores near the A/B transition (compartmental domain boundaries). Grey: Mock; Red: SARS-CoV-2. Compartmental domains were defined with E1 scores using HOMER (findHiCCompartments.pl)(see **Methods**). **c.** Snapshots of inter-chromosomal Hi-C contact matrices between chr17 and chr18 (upper), and intra-chromosomal Hi-C contact matrices within chr18 (lower) in Mock and SARS-CoV-2 infected samples. On the right, differential contacts are shown as log2 fold changes of SARS-CoV-2/Mock. PCA E1 scores were put at sides to show the A or B compartments. Red and black arrowheads respectively point to increased A-B and reduced A-A interactions after SARS-CoV-2 infection.

**Extended Data Figure 4.**
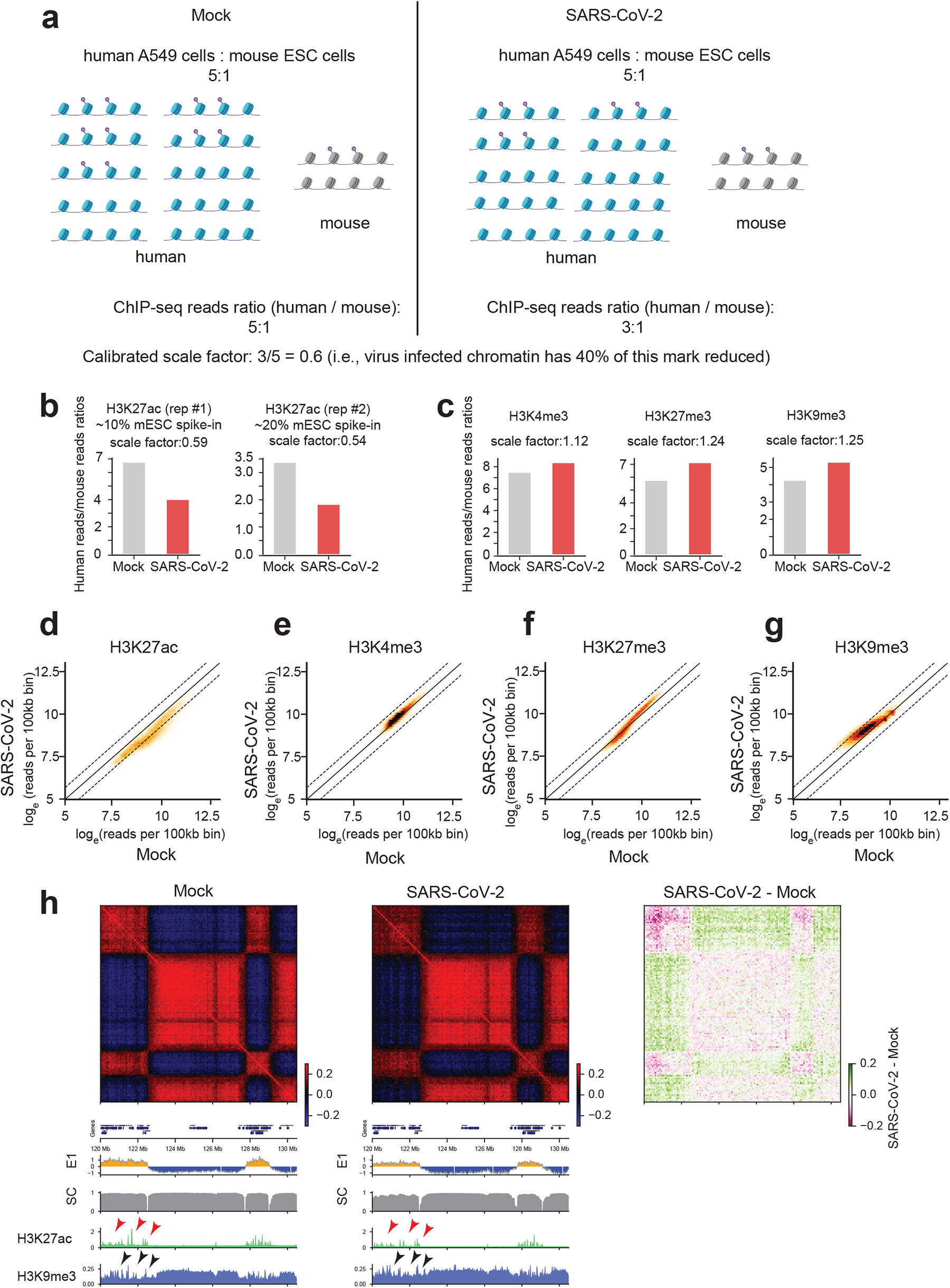
Calibrated ChIP-Seqs demonstrate epigenome reprogramming by SARS-CoV-2 infection. **a.** A diagram illustrating the design of spike-in calibrated ChIP-Seq using mouse ESCs as spike-in for human A549 cells (with or without infection). **b,c.** Barplots showing the human/mouse reads ratio in both Mock and SARS-CoV-2 conditions which permit calibrated ChIP-Seq analyses of H3K27ac, H3K4me3, H3K27me3, and H3K9me3. A scale factor for each histone mark ChIP-seq was denoted above each plot. **d,e,f,g.** Scatter plots show virus-caused genome-wide changes of histone mark ChIP-Seq signals at 100kb bins for H3K27ac, H3K4me3, H3K9me3 and H3K27me3. The x,y-axis are natural logarithmically scaled reads densities from calibrated ChIP-Seq data. Dotted lines denote changes by two folds. **h.** Snapshots of Pearson’s correlation matrices, E1 compartmental scores, sliding correlation scores (SC), as well as ChIP-Seq tracks of H3K27ac and H3K9me3 in Mock or SARS-CoV-2 infected cells. The right side shows the difference of Pearson’s correlation matrices between SARS-CoV-2 and Mock (pink shows decrease). Red arrowheads on top of the H3K27ac peaks show strong reduction of this active mark after infection, which was accompanied by quantitative increase of H3K9me3 signals at the same region (black arrowheads). Accordingly, this entire A compartment shows reduced PCA E1 scores (yellow in the E1 track), showing less compartmental interactions within the same compartment but more interactions with nearby B compartments (see the differential Pearson’s correlation matrices to the right).

**Extended Data Figure 5.**
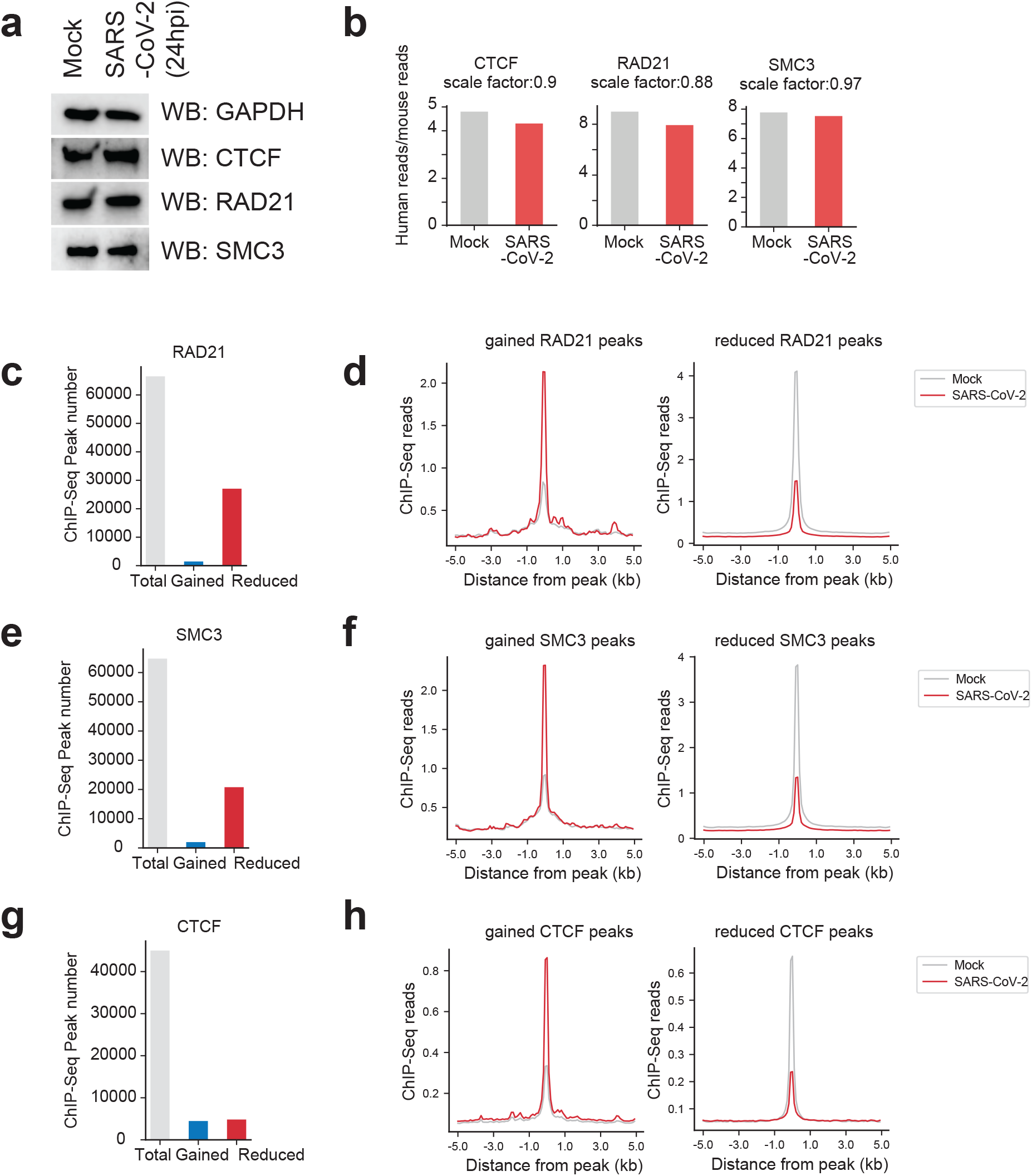
Cohesin depletion specifically from intra-TAD regions after SARS-CoV-2 infection. **a.** Western blots showing the protein abundance of cohesin (RAD21, SMC3) and CTCF in Mock and SARS-CoV-2 infected cells. GAPDH was used as a loading control. **b.** Barplots showing the human/mouse reads ratio in both Mock and SARS-CoV-2 conditions that permit calibrated ChIP-Seqs of CTCF, RAD21, SMC3 and RNA Pol2 (RPB1). These factors were not globally affected by virus infection (so the ratios are comparable in mock and infected conditions). **c.** Barplot showing the number of total, gained or lost RAD21 ChIP-Seq peaks after SARS-CoV-2 infection at 24hpi. **d.** Profile plots showing the signals of RAD21 ChIP-Seq on the gained or lost ChIP-Seq peaks in Mock and SARS-CoV-2 conditions. **e,f,g,h**. Similar to panels c,d, these panels are generated based on calibrated ChIP-Seqs of SMC3 and CTCF respectively.

**Extended Data Figure 6.**
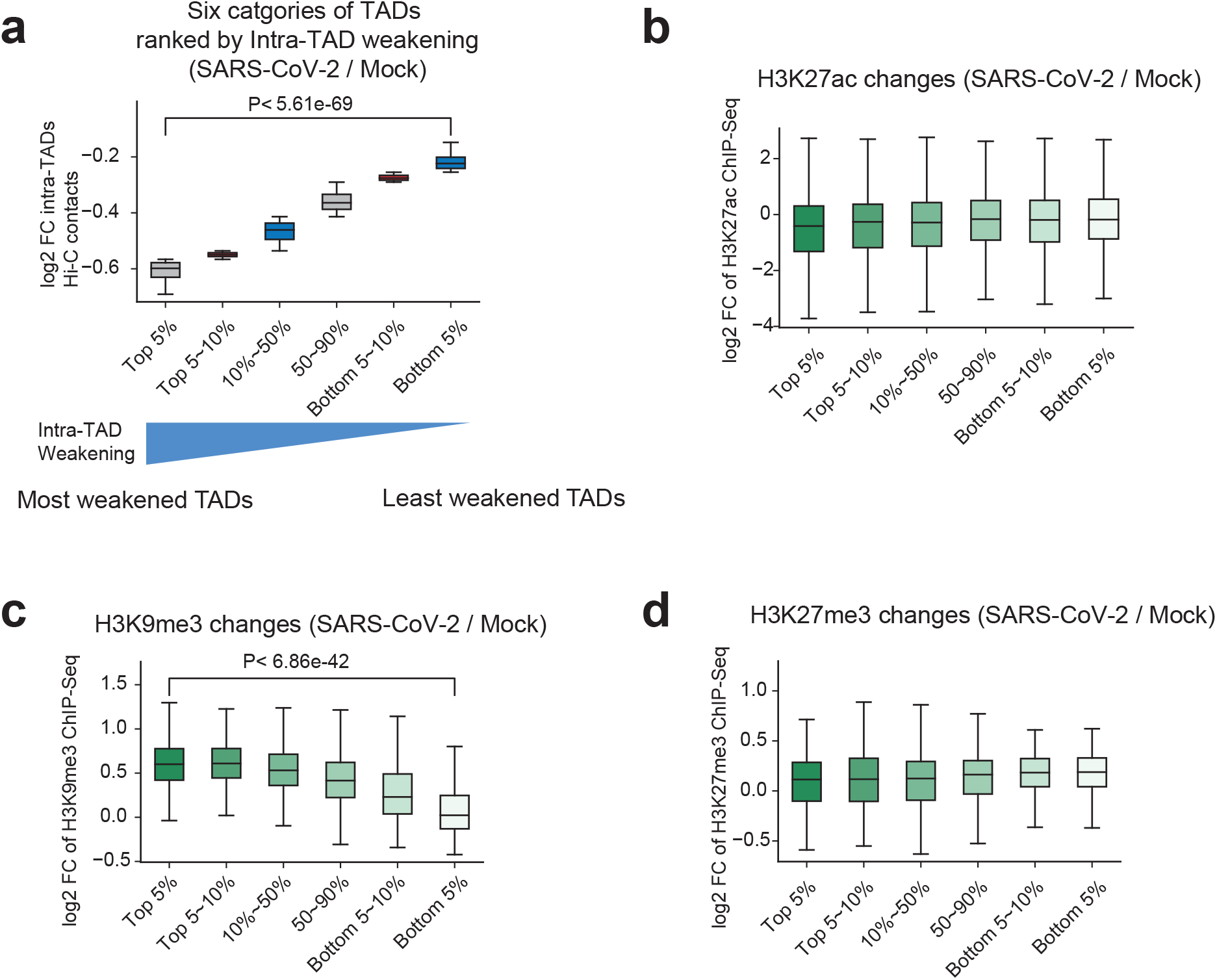
Globally weakened intra-TADs chromatin interactions and the association with altered repressive histone modifications. **a**. A boxplot showing the log2 fold changes of intra-TAD Hi-C contacts for six categories of TADs. All TADs are ranked based on the quantitative changes of intra-TADs interactions and the six categories include the Top 5%, top 5∼10%, 10∼50%, 50∼90%, bottom 5∼10%, and the bottom 5%, respectively. **b,c,d.** Boxplots showing the log2 fold changes of H3K27ac, H3K9me3 and H3K27me3 ChIP-Seq signals in the six categories of TADs as shown in panel a. P-value in panel d: Mann-whitney U test. For all boxplots, the centre lines represent medians; box limits indicate the 25th and 75th percentiles; and whiskers extend 1.5 times the interquartile range (IQR) from the 25th and 75th percentiles.

**Extended Data Figure 7.**
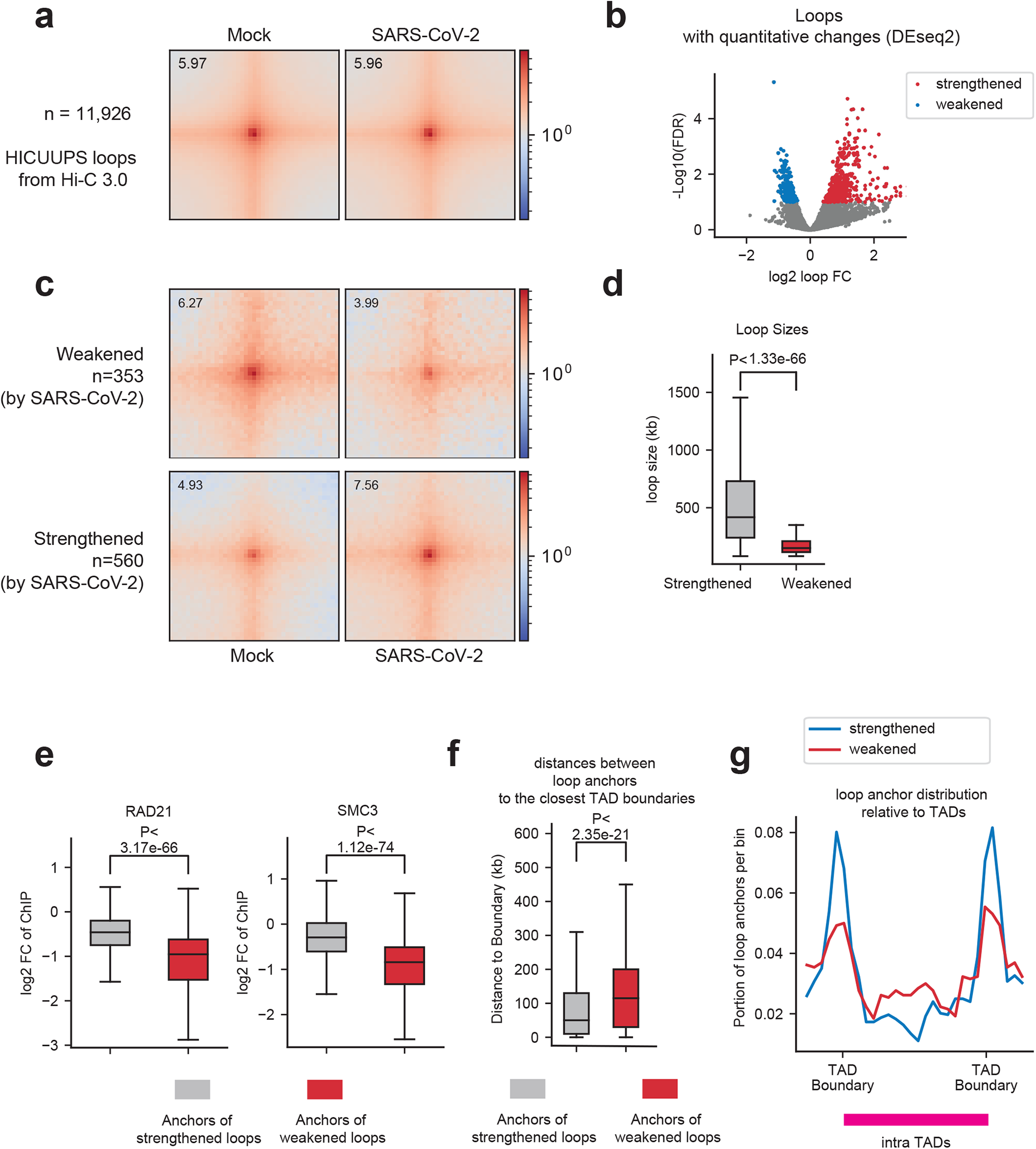
Dot-shaped chromatin loops are largely unaltered after SARS-CoV-2 infection, but a subset of them are changed. **a.** Aggregated peak analysis (APA) shows the strength of all chromatin loops (observed/expected) in Mock (left) and SARS-CoV-2 (right) infected cells for 11,926 dot-shaped loops called by HICCUPS algorithm. **b.** Volcano plot generated by DEseq2 using the two replicates of Hi-C 3.0 that defines quantitatively changed loops after SARS-CoV-2 infection (FDR < 0.1, see **Methods**^25^). **c.** APA plots for the subsets of virus-weakened and strengthened loops. The numbers of such loops are shown. For APA plots in panels a and c, the bin size for plotting the heatmap is 5kb, and the heatmaps show genomic regions +/-100kb surrounding the loop anchors. The numbers on the heatmaps indicate the central pixel values. **d.** A boxplot showing the loop sizes of strengthened and weakened loops. P-values: Mann-whitney U test. **e.** Boxplots showing the virus induced fold changes of cohesin binding (measured by calibrated cohesin ChIP-Seq reads) on the loop anchors of the two groups of loops: those quantitatively strengthened or weakened. P-values: Mann-whitney U test. **f.** Boxplots showing the distances of each loop anchor to its closest TAD boundary for two groups of virus-affected loops: those quantitatively strengthened or weakened. P-values: Mann-whitney U test. **g.** The distribution of loop anchors based on locations relative to its hosting TAD for the two groups of virus-affected loops: those quantitatively strengthened or weakened. Y-axis indicates the percentage of loop anchors in each position relative to hosting TADs. For all boxplots, the centre lines represent medians; box limits indicate the 25th and 75th percentiles; and whiskers extend 1.5 times the interquartile range (IQR) from the 25th and 75th percentiles.

**Extended Data Figure 8.**
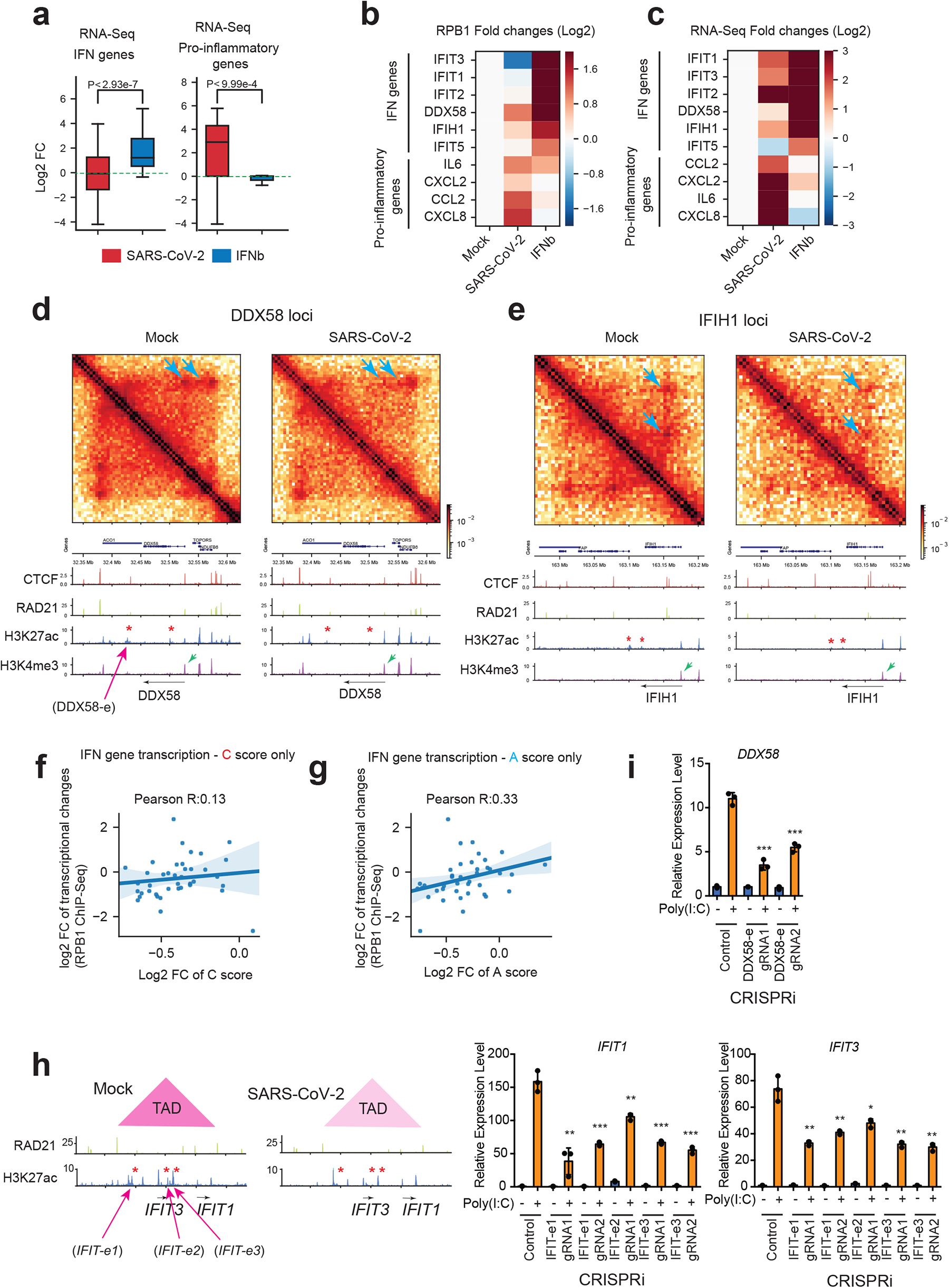
SARS-CoV-2 disrupts chromatin architecture to antagonize the transcriptional activation of interferon response genes. **a.** Boxplots showing the expression deregulation of key interferon response (IFN) and pro-inflammatory genes after SARS-CoV-2 infection or IFN-beta treatment (1000u, 6-hr), as shown by RNA-Seq changes. For boxplots, the centre lines represent medians; box limits indicate the 25th and 75th percentiles; and whiskers extend 1.5 times the interquartile range (IQR) from the 25th and 75th percentiles. P-value: Mann-whitney U test. **b,c.** Heatmaps of select IFN or pro-inflammatory genes showing their fold changes in Pol2 ChIP-Seq or RNA-seq after SARS-CoV-2 infection or IFN-beta treatment (1000u, 6-hr). **d,e.** Snapshots of Hi-C contact matrices and calibrated ChIP-Seq tracks for indicated factors at two key loci coding for virus RNA sensors: *DDX58* (coding for *RIG-I*) and *IFIH1* (coding for *MDA5*). Left: Mock; right: SARS-CoV-2. Blue arrows point to reduced dot-shaped loops. The intra-TAD interactions were weakened throughout these two TADs. Red asterisks show reduced H3K27ac peaks by virus infection. Green arrows show H3K4me3 peaks that are not changed for these genes. **f.** A scatter plot showing a poor correlation between the C score only (from Hi-C contact, x-axis) and the true transcriptional changes of IFN genes by SARS-CoV-2 (y-axis, RPB1 ChIP-Seq). A liner regression fitted line and its 95% confidence interval are also shown. **g.** Similar to panel f, this is a scatter plot showing a poor correlation between the A score only (from enhancer H3K27ac activity, x-axis) and the true transcriptional changes of IFN genes by SARS-CoV-2 (y-axis, RPB1 ChIP-Seq). **h.** (Left) Design of gRNAs/CRISPRi that target the weakened enhancers seen in SARS-CoV-2 infected cells (the red asterisks); this *IFIT* locus is also shown in **Fig. 4b** with more information there. (Right) RT-qPCR results showing that CRISPRi inhibition of the enhancers reduced *IFIT* gene expression in response to poly(I:C), a synthetic viral mimicry. gRNA1 and gRNA2 are two gRNAs targeting the same enhancer (see **Extended Data Table 2**). **i.** Similar to panel h, this is the RT-qPCR result showing that CRISPRi inhibition of the enhancer in *DDX58* locus reduced its response to poly(I:C). The *DDX58* enhancer location for the CRISPRi is shown in panel d. Data in panels h,i show Mean +/-SD from three biological replicates (n=3); p values: two tailed student’s T-test (*, p<0.05; **, p<0.01; ***, p<0.001)

**Extended Data Figure 9.**
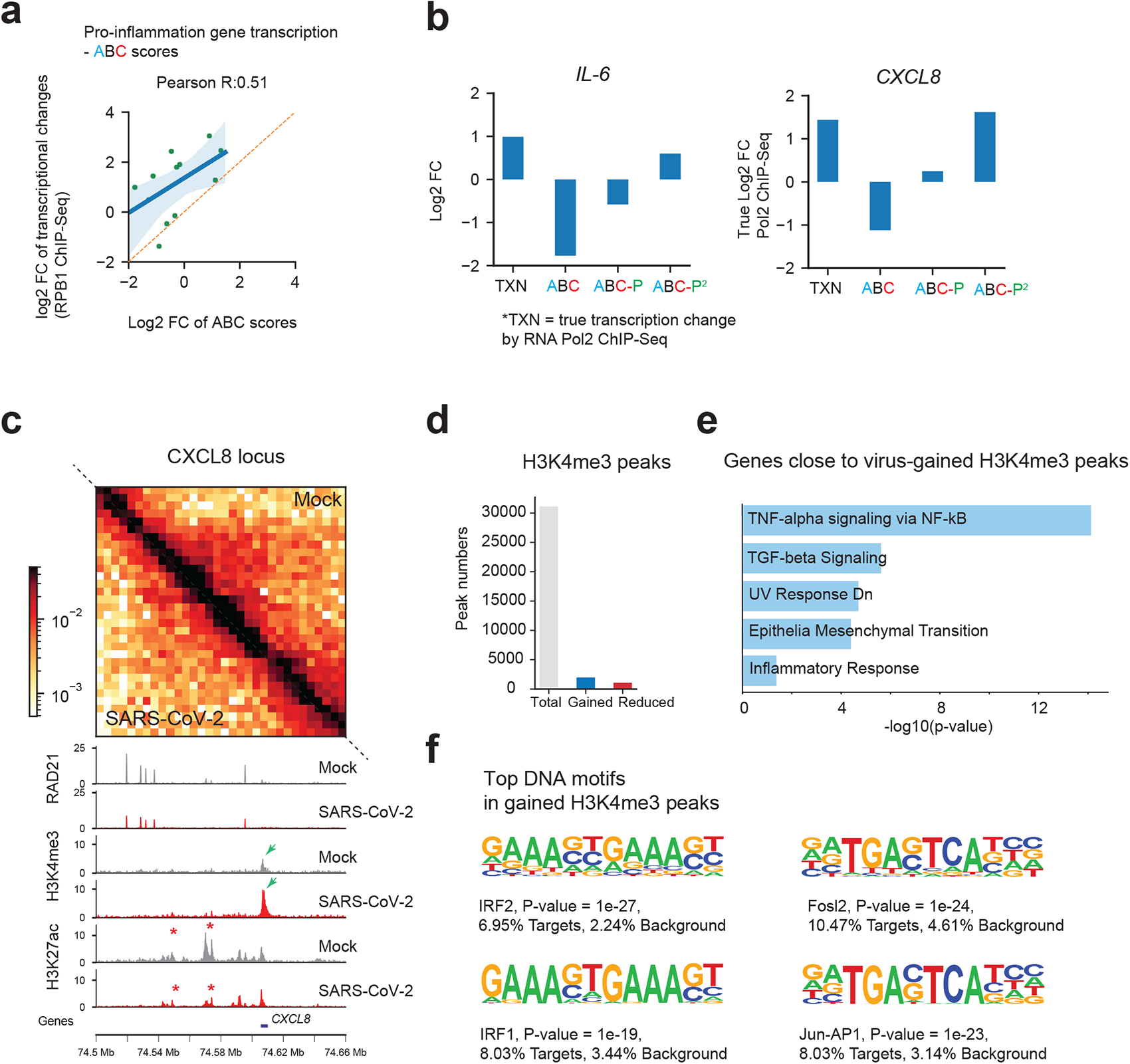
Pro-inflammatory genes were induced by SARS-CoV-2 via augmented promoter activity. **a.** A scatter plot showing good correlation between the ABC score (x-axis) and the true transcriptional changes of pro-inflammatory (PIF) genes by SARS-CoV-2 (y-axis, RPB1 ChIP-Seq). But the true changes (y axis) are much higher than the modeled changes based on ABC scores (x-axis), as all the data points are above the diagonal (also see **Fig. 4f** for revised ABC-P^2^ scores). A liner regression fitted line and its 95% confidence interval are also shown. **b.** Bar graphs showing the fold changes of two key PIF genes, *IL-6* and *CXCL8*, in several conditions: true RPB1 ChIP-Seq fold changes after SARS-CoV-2 infection (TXN); fold changes modeled by ABC score; modeled by ABC algorithm with inclusion of promoter H3K4me3 strength (ABC-P); modeled by ABC algorithm with inclusion of a square of promoter H3K4me3 strength (ABC-P^2^). Promoter strength is required to revise the ABC algorithm to model transcriptional increases of pro-inflammatory genes after SARS-CoV-2 infection. **c.** Snapshots of Hi-C contact matrices and calibrated ChIP-Seq tracks for indicated factors at another key gene loci coding for pathologically critical proinflammatory cytokines in COVID-19 patients: *CXCL8* (a.k.a. *IL-8*). Red asterisks show reduced H3K27ac peaks. Green arrows show increased H3K4me3 peaks at its promoter by SARS-CoV-2 infection. The intra-TAD interactions were weakened throughout the TAD. **d.** Barplot showing the numbers of total, gained or reduced H3K4me3 ChIP-Seq peaks after SARS-CoV-2 infection for 24hpi. **e.** Hallmark signature analysis of genes close to H3K4me3 peaks gained in virus-infected condition show gene signatures associated with TNF-alpha, TGF-beta signaling or inflammatory responses, which are associated with pathological symptoms in COVID-19 patients. **g.** Motif analysis of H3K4me3 peaks increased in SARS-CoV-2 infected cells show that the top ranked motifs are IRF1/2 and Jun/AP1, suggesting their potential roles in transcriptional activation of inflammation genes. Motif analysis was done by HOMER, and the P values and percentages of sites with motifs are shown.

**Extended Data Table 1.** A summary of the Hi-C 3.0, calibrated ChIP-Seq, RNA-Seq datasets generated in this study.

**Extended Data Table 2.** Primers and oligos used in this study.

**Extended Data Table 3.** Gene lists of IFN (interferon response genes) and PIF (pro-inflammatory) genes used in this study.

